# Disrupting Myeloid Persistence and Replenishment Enables Sustained Control of Esophageal Squamous Cell Carcinoma

**DOI:** 10.64898/2026.08.27.747541

**Authors:** Bryan Chee-chad Lung, Alvin Ka-kiu Leung, Songran Liu, Carissa Wing-Yan Wong, Lai Tsz Hong, Ian Yu-hong Wong, Cheryl Chee Heng Lung, Anthony Wing-ip Lo, Ngar-Woon Kam, Josephine Mun-Yee Ko, Wei Dai, Dora Lai-wan Kwong, Simon Law, Pablo Scodeller, Maria Li Lung, Valen Zhuoyou Yu

## Abstract

Responses to macrophage-directed therapy can be transient because tumors preserve myeloid support through complementary persistence and replenishment. In esophageal squamous cell carcinoma (ESCC), CSF1R inhibition reduced established tumor-associated macrophages but was followed by expansion of Ly6C/CCR2-positive monocytic and Ly6G-positive granulocytic populations. Low-dose decitabine preferentially restricted recruited populations while sparing a LYVE1-associated macrophage state, exposing reciprocal pharmacologic blind spots. Combined treatment suppressed both arms and produced sustained control across patient-derived organoid xenograft, orthotopic, and immunocompetent models. Neutrophil depletion reproduced initial regression but not sustained control, indicating that the recruited escape arm extended beyond Ly6G-positive granulocytes. Single-cell profiling mapped these vulnerabilities onto a treatment-resolved myeloid architecture comprising a C1qa-positive TAM continuum, a C1qa-negative Ccr2/Ly6c2-high inflammatory monocytic-like compartment, and a LYVE1/MRC1-positive tissue-supportive macrophage state. Human ESCC contained corresponding macrophage programs and an adverse-outcome-associated LYVE1-rich niche. These findings identify state-aware coverage of complementary myeloid vulnerabilities as a strategy to overcome escape from macrophage-directed therapy.

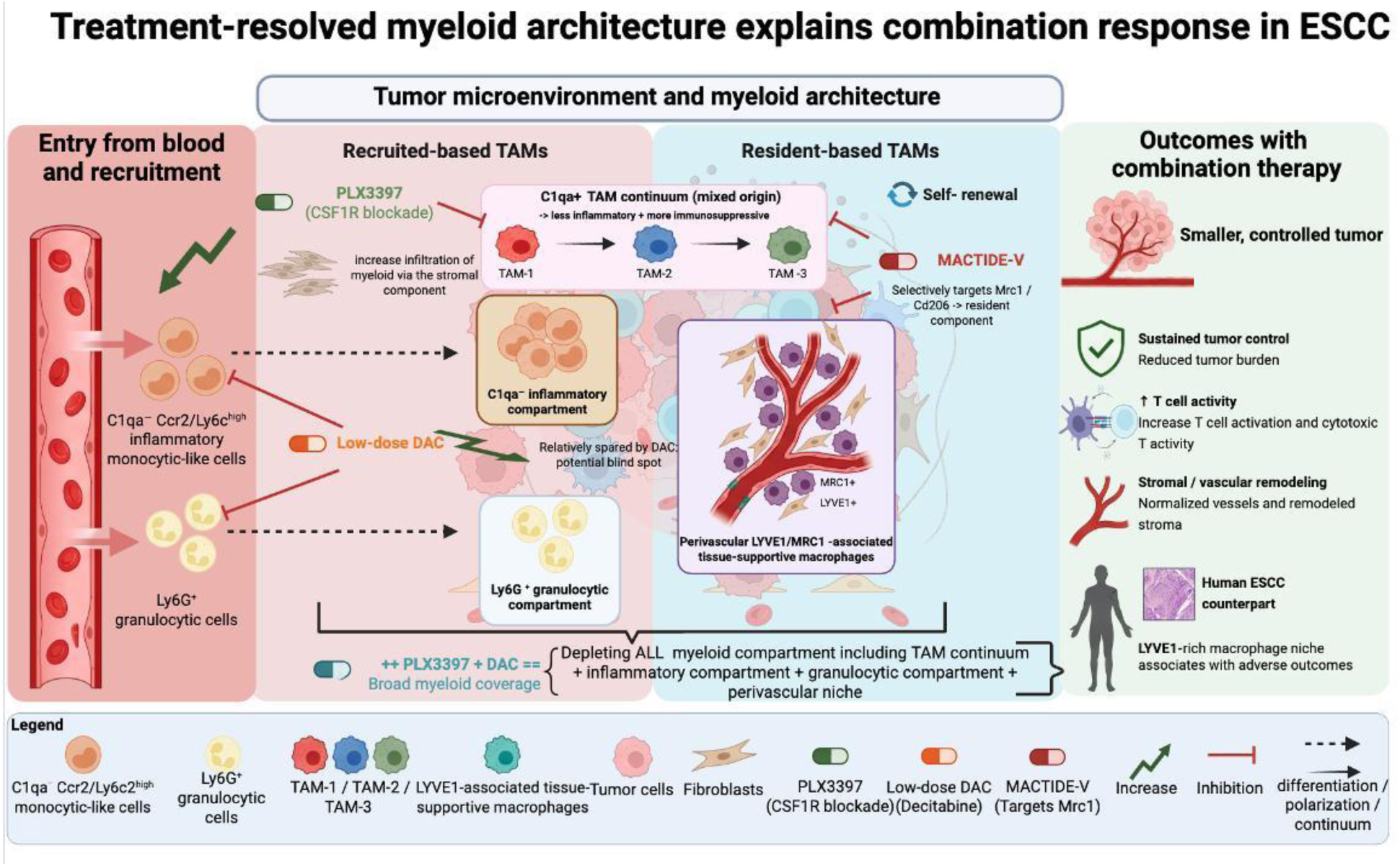

**Statement of Significance:** Macrophage-directed therapy can fail when established tumor-associated macrophage states and recruited myeloid populations remain incompletely covered by a single intervention. In ESCC, jointly suppressing complementary persistence and replenishment compartments converted heterogeneous, incomplete responses into sustained control across models, defining a state-aware myeloid coverage principle for biomarker-guided combination therapy.

## Introduction

Esophageal squamous cell carcinoma (ESCC) remains a lethal malignancy, with 5-year survival of approximately 15% to 20% despite advances in screening, local therapy, and systemic treatment (1–3). Programmed cell death protein 1 (PD-1)-based therapy has improved outcomes for subsets of patients with advanced disease, but durable control remains uncommon, underscoring tumor-microenvironmental programs that sustain resistance (4, 5). Tumor-associated macrophages (TAMs) are abundant components of this ecosystem and can coordinate immunosuppression, angiogenesis, extracellular-matrix remodeling, and treatment resistance (6–8).

The binary M1/M2 framework does not capture the transcriptional, spatial, and functional diversity of tumor myeloid cells (9, 10). Single-cell studies instead resolve antigen-presenting, complement-rich, lipid-handling, tissue-remodeling, and angiogenic macrophage states alongside monocytic and granulocytic populations at different stages of recruitment and differentiation. Yet such atlases often remain descriptive: cluster identity alone does not establish which states persist under therapy, which expand or enter the tumor during escape, or which must be co-targeted. Resident-like and recruited-like compartments may occupy nonredundant niches, creating the possibility that depletion of one compartment permits persistence, expansion, or influx of another (11–13). Because marker expression alone cannot establish developmental origin, these terms require operational rather than lineage-based use. A therapeutically useful macrophage framework must therefore connect state identity to differential treatment vulnerability.

LYVE1-expressing macrophages exemplify a specialized tissue-supportive state. Across nonmalignant and malignant tissues, LYVE1-positive macrophages have been associated with perivascular localization, extracellular-matrix organization, tissue repair, and angiogenesis (14–18). Whether this state forms a treatment-persistent, stromal-supportive niche in ESCC remains unknown.

Colony-stimulating factor 1 receptor (CSF1R) inhibition can reduce TAM abundance, but responses in solid tumors are often incomplete or transient as alternative myeloid and stromal programs persist or emerge (19–24). Low-dose decitabine (DAC), a DNA hypomethylating agent, can reduce suppressive myeloid populations and alter myelopoiesis, although its pharmacology extends to tumor cells, hematopoietic progenitors, stromal cells, and other immune compartments (25–29). We therefore used DAC as a clinically available complementary perturbation—not as a myeloid-specific reagent—to test whether restricting the recruited arm could improve the incomplete control achieved with CSF1R blockade. Rather than assuming either agent was cell type-specific, we asked whether differential treatment sensitivity could resolve states linked to escape and rational combination response. This design addressed a therapeutically consequential question: does incomplete macrophage control reflect persistence of established TAMs, replenishment by recruited cells, or both?

Here, we combined ESCC patient-derived organoid xenograft (PDOX), cell line-derived xenograft (CDX), orthotopic, and immunocompetent models with flow cytometry, immunohistochemistry, bulk transcriptomics, and single-cell RNA sequencing. Across these systems, CSF1R blockade reduced established TAMs but was followed by expansion of recruited myeloid populations, whereas low-dose DAC preferentially restricted the recruited arm while sparing a LYVE1-associated macrophage compartment. Combined treatment suppressed both treatment-defined blind spots, sustained tumor control, and remodeled stromal and immune programs. Single-cell profiling mapped this pharmacologic division onto a treatment-resolved myeloid architecture, and flow cytometry validated its principal vulnerabilities across models. Throughout, resident-like and recruited-like are operational descriptors based on transcriptional profiles, marker expression, flow-cytometric phenotype, and differential treatment sensitivity; definitive lineage tracing was not performed.

## Results

### CSF1R blockade reduces established TAMs but exposes a recruited myeloid escape compartment

To distinguish initial activity from sustained control, we treated mice bearing six independent ESCC PDOX lines with oral pexidartinib (PLX3397; 40 mg/kg daily). PLX3397 slowed tumor growth in all six models, but only two underwent sustained regression; the other four showed partial inhibition or approximately 1 week of tumor stasis before growth resumed (Fig. 1A and B). This broad initial activity but variable durability created an opportunity to distinguish treatment-sensitive TAM states from populations that persisted or expanded during escape.

**Figure 1.**
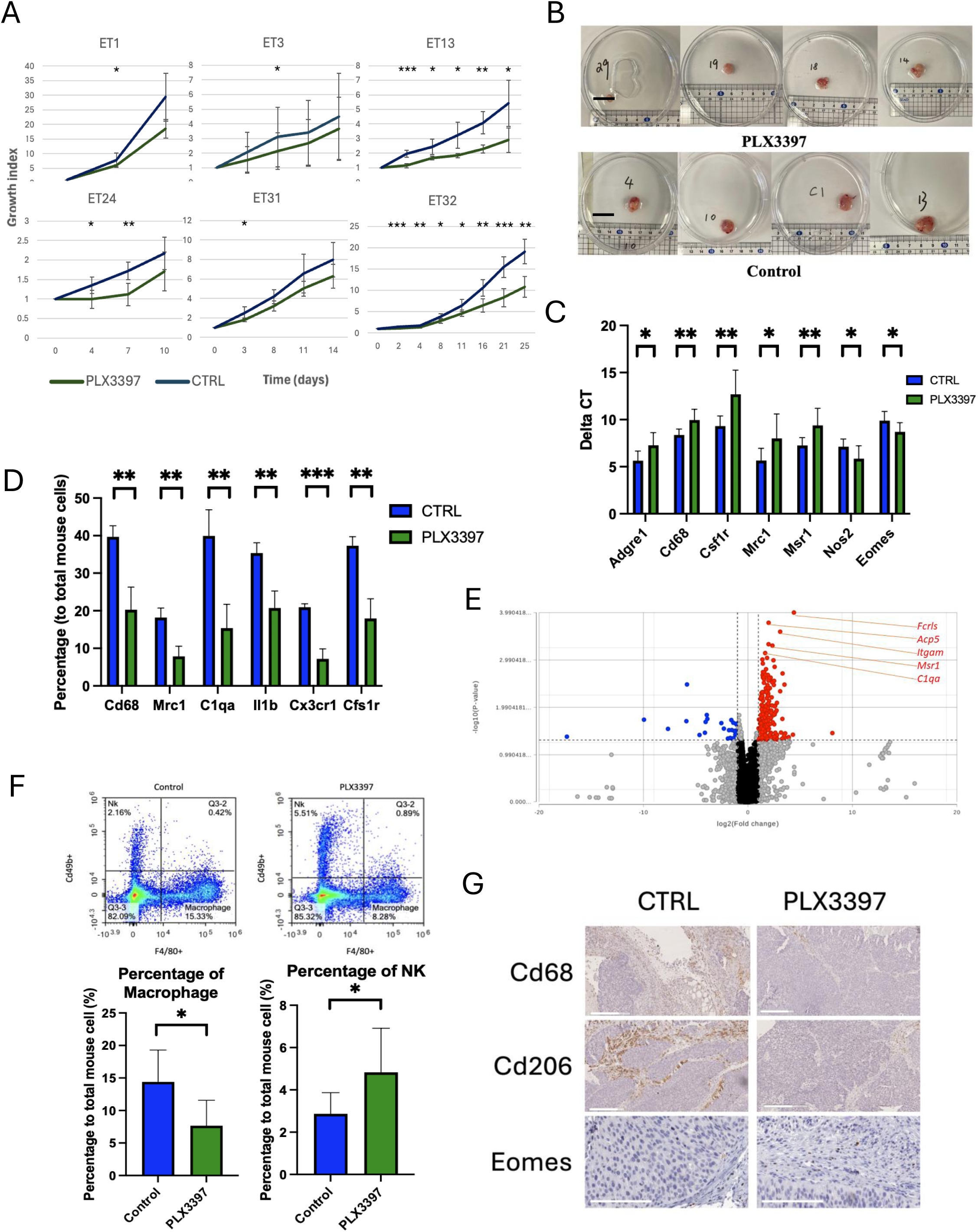
CSF1R blockade reduces established TAMs but does not prevent recruited myeloid replenishment. (**A**) Tumor growth curves of six ESCC PDOX models treated with PLX3397 (40 mg/kg, orally, once daily), illustrating sustained versus partial/transient treatment responses. (**B**) Representative images of PDOX tumors following PLX3397 or solvent control treatment. Scale bar, 20 mm. (**C**) Bulk qPCR analysis of PDOX tumors showing reduced expression of myeloid markers and NK cell–associated genes following PLX3397 treatment. (**D**) Quantification of myeloid cell populations, normalized to total murine cells, from PDOX single-cell RNA sequencing data. (**E**) Bulk transcriptomic analysis of ET1 PDOX tumors demonstrating downregulation of murine myeloid-associated genes following PLX3397 treatment. (**F**) Representative flow cytometry plots (top) and quantification (bottom) of macrophage and NK cell populations after PLX3397 treatment, normalized to total mouse cells. Macrophages were gated as H-2Kd*^+^* F4/80^+^; NK cell H-2Kd^+^ F4/80^-^ Cd49b^+^. (**G**) Representative immunohistochemical staining for Cd68, Cd206, and Eomes in control and PLX3397-treated PDOX tumors. Scale bar, 200 μm. *, P < 0.05; **, P < 0.01; ***, P < 0.001.

PLX3397 reduced established TAM burden across orthogonal assays. Quantitative PCR, bulk and single-cell transcriptomics, flow cytometry, and immunohistochemistry showed lower expression or abundance of F4/80/Adgre1, Cd68, C1qa, Mrc1/CD206, and Msr1 after treatment (Fig. 1C-G). Residual macrophages showed higher Nos2 and lower Mrc1- and Msr1-associated features, indicating altered compartment composition and state rather than proven reprogramming of the same cells. PLX3397-treated tumors also showed increased Eomes expression and greater abundance of CD49b-positive or Eomes-positive natural killer (NK) cells (Fig. 1C, F, and G). The first perturbation therefore defined a PLX3397-sensitive established-TAM compartment while leaving a compositionally distinct, NK-enriched residual microenvironment; NK-cell dependence was not established.

At tumor regrowth, recruited myeloid populations expanded in blood and tumors. Flow cytometry showed increased Ly6C-high monocytic cells and Ly6G-positive granulocytic cells, whereas single-cell analysis showed enrichment of S100a9-, Ly6c2-, and Ccr2-expressing populations (Supplementary Fig. S1A and S1B). Because suppressive function was not directly tested, we refer to Ly6G-positive cells as granulocytes or PMN-MDSC-like cells rather than functionally validated PMN-MDSCs. Their expansion during regrowth, together with the anti-Ly6G intervention below, links the granulocytic component to escape. These findings implicate expansion or influx of recruited myeloid populations—rather than incomplete initial TAM reduction alone—as a limit on PLX3397 durability, consistent with compensatory myeloid responses reported after CSF1R blockade (22, 24).

### Low-dose decitabine restricts recruited myeloid populations while sparing a LYVE1-associated macrophage state

To apply a complementary perturbational axis, we treated ESCC-bearing mice with low-dose 5-aza-2′-deoxycytidine (decitabine; DAC; 1 mg/kg intraperitoneally every 3 days). DAC monotherapy did not significantly inhibit tumor growth (Fig. 2A and B), yet within 1 to 2 weeks markedly reduced circulating and intratumoral Ly6C-high monocytic cells and Ly6G-positive granulocytes, together with Ly6c2-, S100a9-, and Ccr2-expressing populations (Fig. 2C and D; Supplementary Fig. S1A). Circulating lymphocyte populations were comparatively preserved (Supplementary Fig. S1B). Under these models and dosing conditions, the dominant observed myeloid effect of DAC was preferential restriction of the recruited arm, not absolute selectivity for a single lineage.

**Figure 2.**
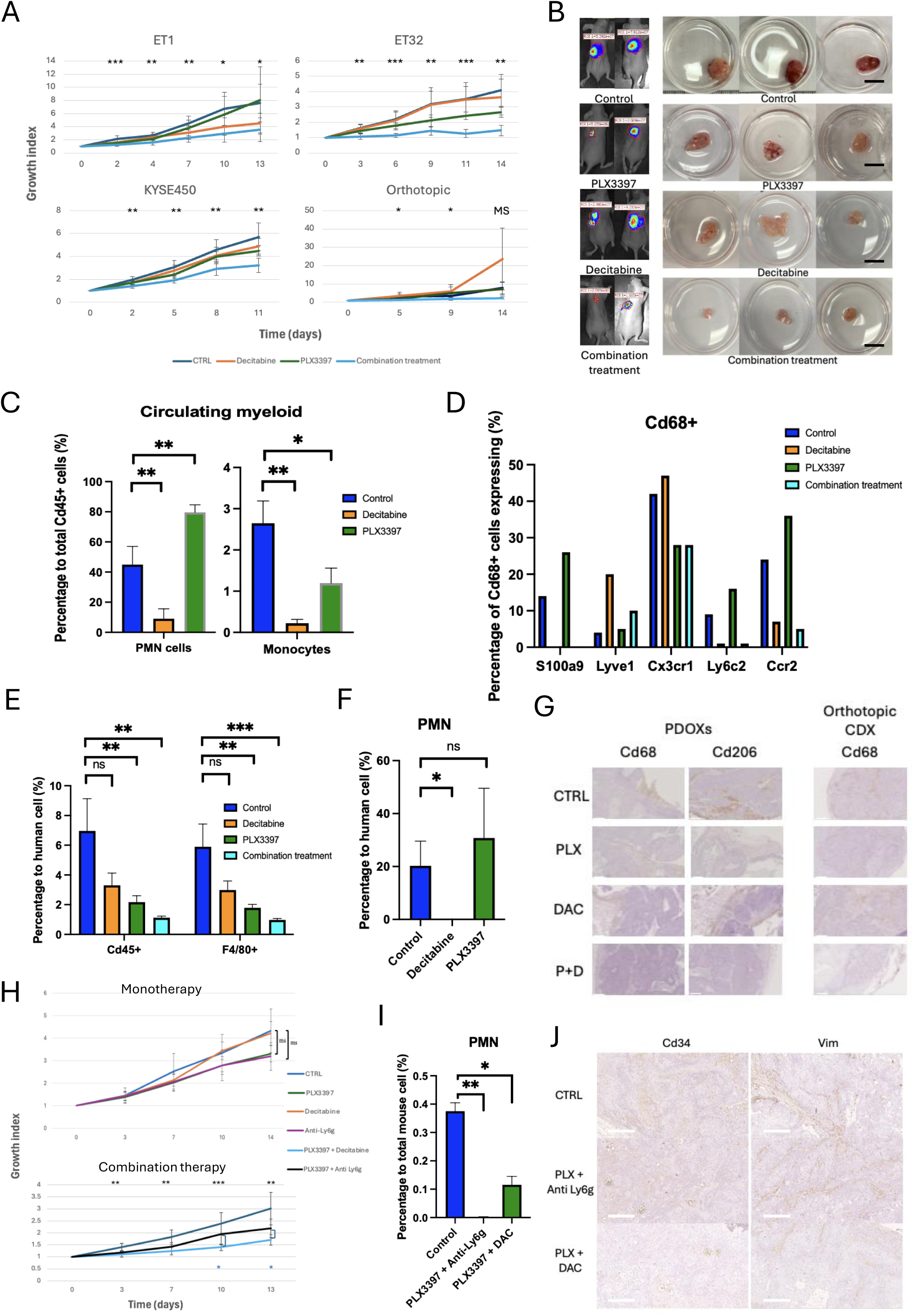
Complementary targeting of established TAMs and recruited myeloid cells produces durable control of ESCC. (**A**) Tumor growth curves of ESCC models treated with vehicle, PLX3397, decitabine (DAC), or the combination. Therapeutic responses are shown across PDOX (ET1 and ET32), subcutaneous CDX (KYSE450), and orthotopic CDX (KYSE450) models. (**B**) Representative images of orthotopic CDX (left) and PDOX (right) tumors following the indicated treatments. Scale bar, 20 mm. (**C**) Representative flow cytometry plots (top) and quantification (bottom) of circulating monocytes and polymorphonuclear (PMN) cells following vehicle, PLX3397, or DAC treatment. Monocytes were gated as H-2Kd⁺Cd45⁺Ly6g⁻Cd49b⁻F4/80⁻Cd11c⁻Ly6c⁺ cells, and PMN cells as H-2Kd⁺Cd45⁺Cd49b⁻Ly6g⁺ cells. (**D**) Percentage of CD68⁺ myeloid cells expressing the indicated genes across treatment groups, determined by single-cell RNA sequencing. (**E**) Representative flow cytometry plots (top) and quantification (bottom) of total CD45⁺ immune cells and F4/80⁺ tumor-associated macrophages (TAMs), normalized to human EPCAM⁺ tumor cells. Immune cells were gated as H-2Kd⁺CD45⁺EPCAM⁻ cells, and TAMs as H-2Kd⁺Cd45⁺EPCAM⁻F4/80⁺ cells. (**F**) Flow cytometry quantification of tumor-infiltrating PMN cells, normalized to human EPCAM⁺ tumor cells. PMN cells were gated as H-2Kd⁺Cd45⁺Cd49b⁻Ly6g⁺ cells. (**G**) Representative immunohistochemical staining for Lyve1 in PDOX and orthotopic CDX tumors across treatment groups. Scale bar, 200 μm. (**H**) Tumor growth curves comparing vehicle, PLX3397 and anti-Ly6g, monotherapy (top) and vehicle, PLX3397 plus anti-Ly6G, and PLX3397 plus DAC treatment (bottom). (**I**) Representative flow cytometry plots (top) and quantification (bottom) of tumor-infiltrating PMN cells following the indicated combination therapies. PMN cells were gated as H-2Kd⁺Cd45⁺Cd49b⁻Ly6g⁺ cells. (**J**) Representative immunohistochemical staining for Cd34 (left) and Vim (right), showing endothelial cell and fibroblast abundance in PDOX tumors following PLX3397 plus anti-Ly6g or PLX3397 plus DAC treatment. Scale bar, 200 μm. *, P < 0.05; **, P < 0.01; ***, P < 0.001; MS, marginal significance (P < 0.1).

By contrast, DAC minimally changed the overall frequency of intratumoral F4/80-positive macrophages in PDOX and subcutaneous CDX models (Fig. 2E). Cx3cr1-expressing macrophage states and a LYVE1-associated compartment became proportionally enriched as DAC-sensitive populations declined (Fig. 2D). Flow cytometry and histologic analyses supported persistence of this compartment, reducing the likelihood that the single-cell pattern reflected relative enrichment alone (Fig. 2E-G). DAC did not appear to induce a LYVE1 state de novo; rather, this second perturbation distinguished DAC-sensitive recruited cells from a comparatively spared LYVE1-associated macrophage state. Because DAC is pharmacologically pleiotropic, these results define differential susceptibility in this setting rather than a recruited-myeloid-specific drug effect.

### Combined CSF1R blockade and low-dose decitabine produce sustained ESCC control across models

Having resolved reciprocal monotherapy blind spots, we asked whether combined PLX3397 and DAC could suppress both. The combination produced greater regression or growth inhibition than either monotherapy across the tested ESCC models, including orthotopic tumors (Fig. 2A and B). The combination remained active in PDOX lines with limited monotherapy responsiveness and maintained control through the study period or until a prespecified humane endpoint, providing the operational definition of sustained suppression used in these experiments. Thus, joint treatment converted heterogeneous, incomplete monotherapy responses into sustained cross-model control.

Flow cytometry revealed a marked reduction in CD45-positive leukocytes, near-complete depletion of the major detectable F4/80-positive TAM populations, and loss of Ly6G-positive cells after combination treatment (Fig. 2E and F). Immunohistochemistry likewise showed pronounced reductions in F4/80/CD68-positive macrophage populations (Fig. 2G). Because these xenograft studies used nude hosts, substantial tumor control occurred in the absence of conventional T-cell immunity; this does not exclude contributions from innate immune, stromal, hematopoietic, or tumor-cell effects of either agent.

Combination-treated mice maintained stable body weight (Supplementary Fig. S2A). In vitro exposure of ESCC organoids to PLX3397, DAC, or the combination did not reveal combination-enhanced direct cytotoxicity (Supplementary Fig. S2B). This result supports a major contribution from the in vivo microenvironment but does not exclude low-dose DAC effects on tumor-cell immunogenicity or other nonmyeloid programs. The in vivo data therefore demonstrate cooperative efficacy exceeding either monotherapy, without establishing formal pharmacologic synergy.

### Neutrophil depletion reproduces initial regression but not sustained control

To determine whether Ly6G-positive granulocytes alone represented the recruited compartment responsible for escape, we compared PLX3397 plus anti-Ly6G with PLX3397 plus DAC. Anti-Ly6G monotherapy had minimal antitumor activity, similar to PLX3397 alone (Fig. 2H). PLX3397 plus anti-Ly6G induced rapid regression comparable with the initial response to PLX3397 plus DAC, and both regimens markedly reduced intratumoral Ly6G-positive cells (Fig. 2H and I). These intervention data establish a contribution from the granulocytic branch after TAM depletion.

Despite this initial response, tumors treated with PLX3397 plus anti-Ly6G eventually regrew, whereas PLX3397 plus DAC maintained regression through the study period (Fig. 2H). Tumors in the anti-Ly6G combination group retained greater fibroblast and endothelial abundance than tumors treated with PLX3397 plus DAC (Fig. 2J). Mice receiving PLX3397 plus anti-Ly6G also developed greater weight loss and cachexia, whereas the DAC combination was better tolerated (Supplementary Fig. S2C); systemic inflammation was not directly measured. The later divergence indicates that the granulocytic branch is important but insufficient to explain the full recruited escape arm, implicating broader monocytic and granulocytic state coverage.

### Single-cell profiling maps reciprocal vulnerabilities onto treatment-resolved myeloid states

To resolve the cellular architecture underlying these differential vulnerabilities, we performed single-cell RNA sequencing in two PDOX models with different PLX3397 responses. ET32 tumors treated with vehicle, PLX3397, DAC, or the combination were profiled with three biological replicates per group; ET1 vehicle- and PLX3397-treated tumors were profiled with two biological replicates per group. Approximately 106,000 high-quality transcriptomes were recovered, including approximately 76,000 cells from ET32 and 30,000 from ET1. Alignment to a combined human-mouse reference separated 50,358 EPCAM-positive human ESCC cells from 55,914 murine microenvironmental cells (Fig. 3A; Supplementary Fig. S3A). The murine compartment included cancer-associated fibroblasts (CAFs), pericytes, endothelial cells, mononuclear myeloid cells, NK/innate lymphoid cells, and granulocytes.

**Figure 3.**
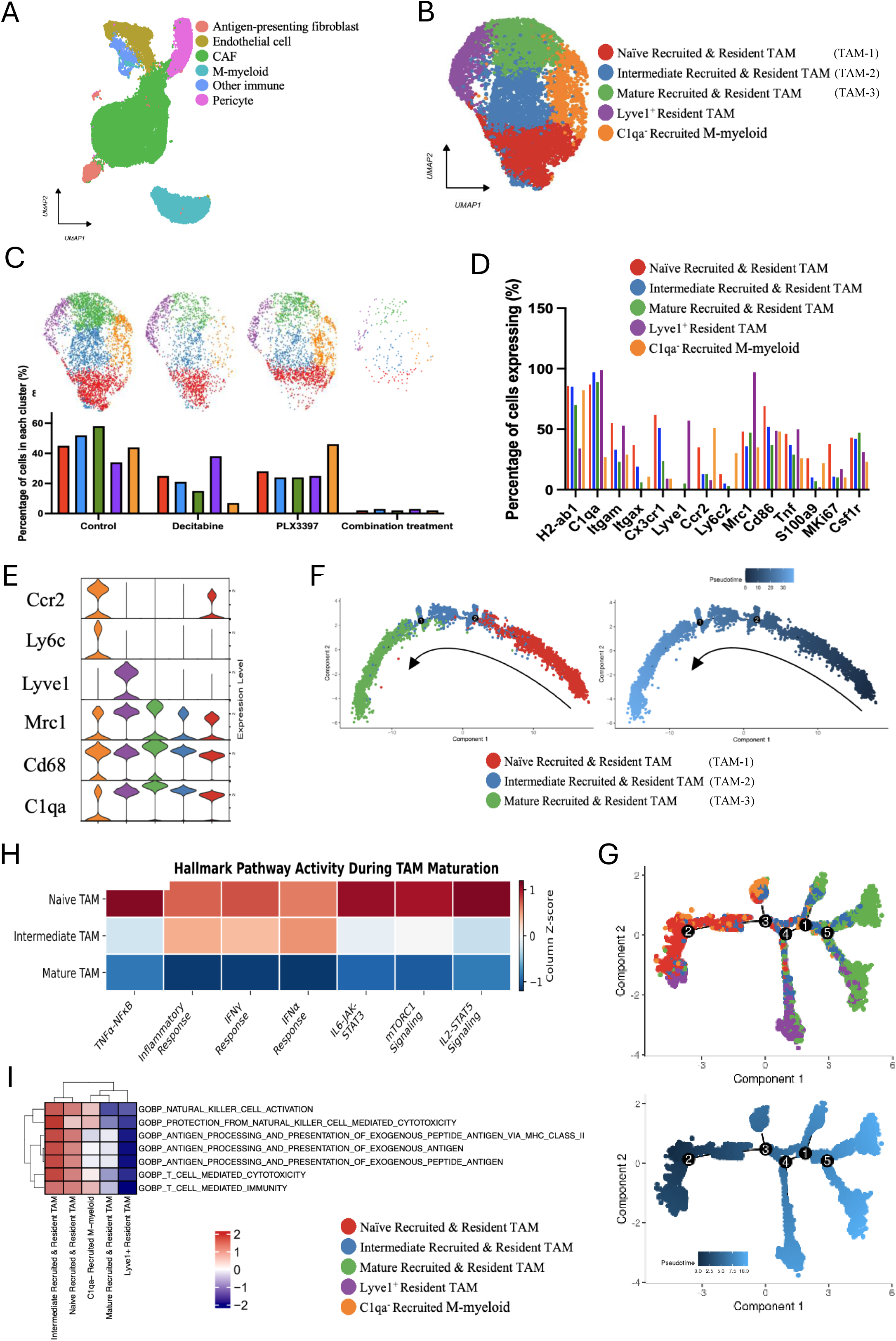
Single-cell profiling resolves complementary resident and recruited myeloid compartments underlying monotherapy escape. (**A**) UMAP visualization of the murine tumor microenvironment following separation of human EPCAM⁺ tumor cells and murine stromal/immune cells. (**B**) Re-clustering of 8,460 CD68^+^ mononuclear myeloid cells identified five distinct M-myeloid subpopulations. (**C**) UMAP plots showing the distribution of M-myeloid cells across treatment groups (top) and the relative abundance of each subpopulation under the indicated treatments (bottom). (**D**) Percentage of cells expressing the indicated myeloid related markers across the five M-myeloid subpopulations. (**E**) Violin plot showing the expression of defining marker genes for each M-myeloid subpopulation. size indicates the percentage of expressing cells, and color denotes average expression level. (**F**) Monocle pseudotime analysis of TAM-1, TAM-2, and TAM-3 demonstrating a continuous differentiation trajectory from inflammatory naïve TAMs through intermediate TAMs to mature TAMs. (**G**) Pseudotime projection of all five M-myeloid subpopulations, illustrating the relationship between different macrophage states. (**H**) Hallmark pathway enrichment analysis across the TAM differentiation trajectory (TAM-1 to TAM-3). (**I**) Gene Ontology Biological Process (GOBP) enrichment analysis of the five M-myeloid subpopulations, highlighting distinct functional programs associated with each macrophage state.

Unsupervised reclustering of 8,460 Cd68-high mononuclear myeloid cells resolved five transcriptionally distinct populations (Fig. 3B and C). Three C1qa-positive populations formed a continuum that we designated TAM-1, TAM-2, and TAM-3. TAM-1 retained the strongest inflammatory and antigen-presentation features, including relatively higher Ly6c2, Ccr2, Cd86, Tnf, and H2-Ab1 expression, and contained a small Mki67-positive fraction. TAM-2 occupied an intermediate transcriptional position, whereas TAM-3 displayed the weakest inflammatory and antigen-presentation programs. Two populations remained outside this core continuum: a LYVE1-positive, Mrc1-high, C1qa-positive tissue-supportive macrophage state and a C1qa-negative inflammatory monocytic-like population (M-myeloid; Ly6c2/Ccr2-high and Cx3cr1/Lyve1-low) (Fig. 3B-E). Together, these populations defined a macrophage-centered myeloid architecture without implying fixed lineage identities.

Treatment response mapped directly onto this architecture. PLX3397 reduced the C1qa-positive TAM continuum, whereas the relative abundance of the C1qa-negative inflammatory monocytic-like population increased (Fig. 3C). This population expressed the lowest Csf1r among the five states, consistent with reduced susceptibility to CSF1R blockade (Fig. 3D). Conversely, DAC nearly eliminated the C1qa-negative population but left the LYVE1-positive macrophage state proportionally enriched. Across the three combination-treated ET32 tumors, only 192 mononuclear myeloid cells were recovered, representing approximately 2% of the profiled mononuclear myeloid compartment (Fig. 3C). Flow cytometry and immunohistochemistry supported the same directional changes, although tumor shrinkage, dissociation, viable-tissue sampling, and differential cell recovery may influence apparent single-cell abundance.

Pseudotime analysis was restricted to TAM-1, TAM-2, and TAM-3 rather than forcing all five populations into a single developmental hierarchy (Fig. 3F and G; Supplementary Fig. S3B). Within the C1qa-positive continuum, progression toward TAM-3 was accompanied by attenuation of interferon-alpha/gamma, inflammatory-response, antigen-presentation, and phagocytosis-associated programs (Fig. 3H and I). C1qa was broadly expressed in differentiated TAM states but absent from the Ccr2/Ly6c2-high population, consistent with acquisition of C1q during macrophage differentiation (30). Only a subset of C1qa-negative cells showed detectable S100a9 at single-cell resolution, indicating biological heterogeneity while leaving transcript dropout as a possible contributor. The resulting architecture therefore comprised a central C1qa-positive TAM-conditioning continuum together with distinct recruited inflammatory monocytic-like and LYVE1-associated tissue-supportive states, rather than a single linear hierarchy.

### Flow cytometry validates complementary treatment-resolved vulnerabilities across ESCC models

To determine whether the single-cell-defined vulnerabilities extended beyond the profiled tumors, we translated their surface-marker features into an operational flow-cytometric classification. Across the examined models, marked depletion of total macrophages was consistently observed only after combined PLX3397 and DAC, whereas DAC-containing regimens strongly reduced Ly6G-positive cells (Fig. 4A; Supplementary Fig. S4A).

**Figure 4.**
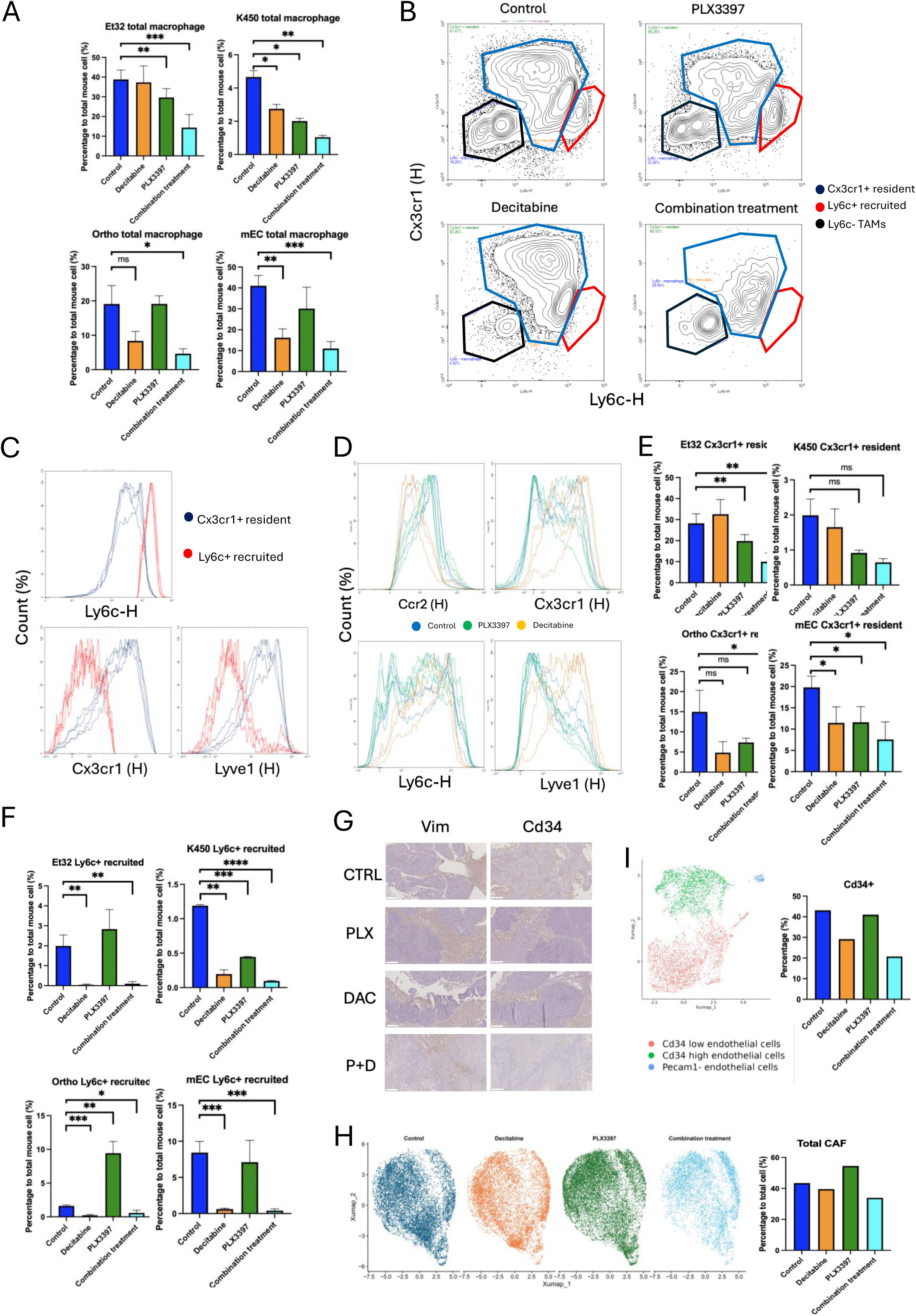
Dual myeloid targeting remodels the tumor microenvironment through coordinated depletion of resident and recruited macrophages. (**A**) Flow cytometric quantification of total TAMs across treatment groups, normalized to total murine cells. TAMs were gated as H-2Kd⁺Cd45⁺EPCAM⁻F4/80⁺ cells. (**B**) Representative flow cytometry plots demonstrating the gating strategy (Cx3cr1 x Ly6c) used to distinguish Ly6c^hi^Cx3cr1^low^ recruited-like and Ly6c^low^Cx3cr1^hi^ resident-like macrophage-enriched compartments across treatment groups. (**C**) Expression of Ly6c, Cx3cr1, and Lyve1 within Ly6C⁺ recruited-like and CX3CR1⁺ resident-like TAM compartments, validating the operational definition of recruited and resident macrophage populations. (**D**) Expression of recruited-like markers (Ly6c2 and Ccr2) and resident-like markers (Cx3cr1 and Lyve1) in TAMs following vehicle, PLX3397, or DAC treatment. (**E**) Flow cytometric quantification of Ly6c⁺ recruited-like TAMs across all PDOX, CDX, and orthotopic models following the indicated treatments. (**F**) Flow cytometric quantification of Cx3cr1⁺ resident-like TAMs across all PDOX, CDX, and orthotopic models following the indicated treatments. (**G**) Representative immunohistochemical staining for Vim (left) and Cd34 (right), demonstrating that only combined PLX3397 and DAC treatment markedly reduced stromal fibroblast and endothelial cell abundance. Scale bar, 200 μm. (**H**) Single-cell RNA sequencing showing total cancer-associated fibroblast (CAF) abundance (top) and the relative distribution of CAFs across treatment groups (bottom) in ET32 PDOX tumors. (**I**) UMAP visualization of endothelial cells (left) and quantification of CD34⁺ endothelial cells (right) across treatment groups in ET32 PDOX tumors. *, P < 0.05; **, P < 0.01; ***, P < 0.001.

Ly6C and CX3CR1 separated macrophage-enriched compartments with complementary treatment sensitivity. Ly6C-high/CX3CR1-low cells were preferentially reduced by DAC, whereas Ly6C-low/CX3CR1-high cells were comparatively spared (Fig. 4B and C). LYVE1 provided an additional readout of the spared macrophage state, although it did not map perfectly onto the Ly6C/CX3CR1 gate. Consistent with the single-cell data, macrophages remaining after PLX3397 showed higher Ly6C and CCR2, whereas those remaining after DAC showed higher CX3CR1 and LYVE1 (Fig. 4D). We therefore use these gates as recruited-like and resident-like operational compartments, not as definitive lineage assignments.

Using this framework, DAC preferentially reduced the recruited-like macrophage-enriched compartment in most tested models, whereas the resident-like compartment was comparatively spared; PLX3397 showed variable depletion of TAM states across models (Fig. 4E and F). Only combined PLX3397 and DAC consistently reduced both compartments across treatment settings, providing an orthogonal explanation for the superior antitumor activity (Fig. 2A). Because baseline immune infiltration and implantation sites differed among models, populations were normalized to total murine cells for cross-group comparison. The reproducible partition of treatment sensitivity across models supports the treatment-resolved architecture while remaining agnostic about developmental origin.

### Broad suppression of treatment-resolved myeloid states coincides with stromal, vascular, and immune remodeling

We next asked whether suppression of both major treatment-defined myeloid arms coincided with broader remodeling of the ESCC microenvironment. Vimentin immunohistochemistry and single-cell analysis showed lower CAF abundance after combined PLX3397 and DAC, accompanied by reduced Acta2 and Fap expression (Fig. 4G and H; Supplementary Fig. S4B). PLX3397 monotherapy was associated with increased CAF abundance in the profiled tumors (Fig. 4H), consistent with compensatory macrophage-fibroblast interactions described in other settings (22, 31).

Endothelial abundance was also reduced after combination treatment. The CD34-positive subset comprised approximately 20% of endothelial cells in combination-treated tumors compared with 43% in controls, and CD34 immunostaining showed lower microvessel density (Fig. 4G and H). Because the combination also reduced the LYVE1-associated angiogenic macrophage state, these findings are consistent with disruption of macrophage-supported vascular niches (14). Tumor shrinkage, viable-tissue sampling, and treatment-related recovery differences may also contribute and preclude definitive causal attribution.

Gene-set enrichment analysis showed increased antigen-processing and presentation programs in human ESCC cells and CAFs after combination treatment; T-cell activation and lymphocyte-mediated immunity signatures were also enriched (Supplementary Fig. S4C and S4D). Broad suppression of the treatment-resolved myeloid states therefore coincided with reduced fibroblast and endothelial abundance and increased antigen-presentation and lymphoid programs. These data do not establish direct stromal causality or adaptive immune dependence.

### Human ESCC preserves corresponding macrophage programs and an adverse LYVE1-associated niche

To determine whether the treatment-linked macrophage states had counterparts in human disease, we reanalyzed a publicly available ESCC single-cell RNA-sequencing dataset using a workflow aligned with the murine analysis (32; Supplementary Fig. S5A and S5B). After exclusion of monocytes and dendritic cells, unsupervised clustering identified five macrophage populations (Fig. 5A). Cross-species comparison linked human clusters to murine TAM-1, TAM-3, and the distinct LYVE1-associated state. Human TAM-1-like cells expressed higher antigen-presentation genes, including HLA-DMB, HLA-C, and HLA-DQA1; cell-cycle genes, including MKI67 and STMN1; and inflammatory monocytic genes, including S100A8 and S100A9. TAM-3-like cells showed lower inflammatory and immune-activating pathway activity, whereas a distinct LYVE1-associated cluster expressed LYVE1, MRC1, and APOE and was enriched for angiogenesis-related features (Fig. 5B and C; Supplementary Fig. S5C). These data support the presence of a corresponding LYVE1-associated macrophage state across species.

**Figure 5.**
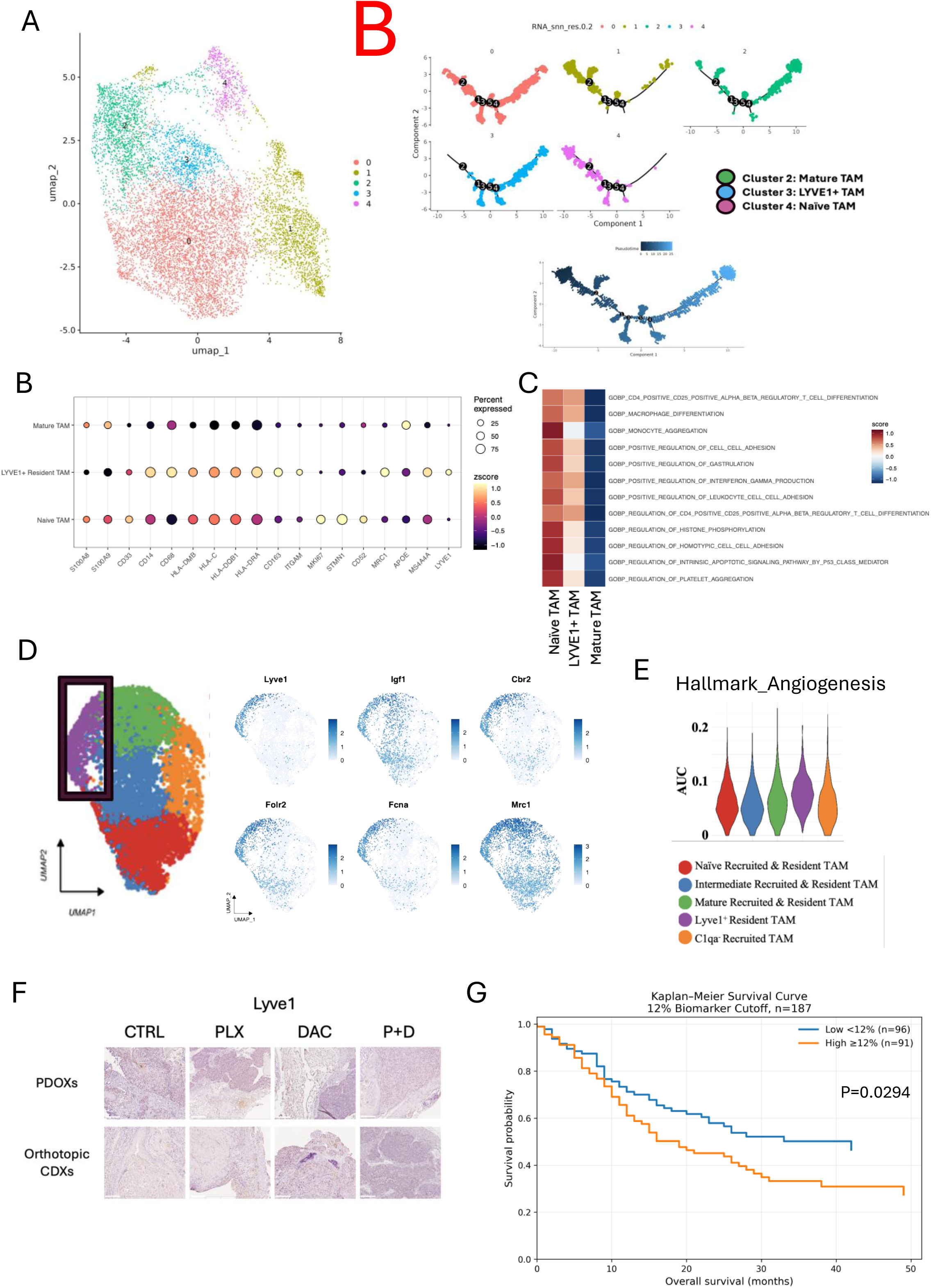
A conserved LYVE1-associated resident macrophage program predicts poor outcome in human ESCC. (**A**) Unsupervised clustering (top) and pseudotime trajectory analysis (bottom) of macrophages from a public single-cell RNA sequencing dataset comprising 60 human ESCC tumors, identifying naïve, mature, and LYVE1-associated macrophage states. (**B**) Dot plot showing the expression of defining macrophage marker genes across human ESCC macrophage subpopulations. Dot size indicates the percentage of expressing cells, and color denotes average expression level. (**C**) Gene Ontology Biological Process (GOBP) enrichment analysis comparing naïve, mature, and LYVE1-associated macrophage states in human ESCC. (**D**) Reference UMAP of murine M-myeloid cells highlighting the Lyve1-associated resident macrophage population (top) and feature plots showing the expression of Lyve1-associated genes across murine M-myeloid cells (bottom). (**E**) Hallmark pathway enrichment analysis demonstrating preferential activation of angiogenesis in the murine Lyve1⁺Folr2⁺Mrc1⁺ resident-like macrophage state. (**F**) Representative Lyve1 immunohistochemical staining in subcutaneous PDOX and orthotopic CDX tumors following the indicated treatments. Orthotopic tumors exhibited greater LYVE1⁺ macrophage abundance, whereas only combined PLX3397 and decitabine (DAC) treatment effectively depleted this population. Scale bar, 200 μm. (**G**) Kaplan–Meier analysis of overall survival in an ESCC tissue microarray cohort stratified by a 12% LYVE1-positivity cutoff.

In murine ESCC, the LYVE1-positive macrophage state preferentially expressed Lyve1, Folr2, Cbr2, Fcna, Mrc1, and Igf1 and was enriched for tissue-remodeling and angiogenic programs (Figs. 3D, 3E, 5D, and 5E). The state lacked Ly6c2 and Ccr2, contained a small Mki67-positive fraction, and remained transcriptionally distinct from both the C1qa-positive TAM continuum and the C1qa-negative inflammatory monocytic-like state. Together with its relative preservation following DAC, these features are consistent with a resident-like, vascular-associated macrophage state distinct from DAC-sensitive monocytic populations, but do not establish developmental origin. Tumor LYVE1 staining was most strongly reduced by combined PLX3397 and DAC in both subcutaneous and orthotopic models (Fig. 5F).

We next evaluated LYVE1 staining in two ESCC tissue microarrays (HEso-Squ172Sur-01 and HEso-Squ180Sur-03) comprising 187 patients. Using an outcome-optimized cutoff of 12% LYVE1 positivity (Supplementary Fig. S6A and S6B), LYVE1-high tissue was associated with poorer overall survival than LYVE1-low tissue (median, 15 vs. 21 months; log-rank P = 0.0294; Fig. 5G; Table 1). High LYVE1 staining was associated with increased mortality in univariable analysis (HR, 1.53; 95% CI, 1.04-2.27; P = 0.0328) and after adjustment for age and sex (HR, 1.66; 95% CI, 1.12-2.47; P = 0.0119; Table 1; Supplementary Table S1). Because LYVE1 is not macrophage-specific, macrophage colocalization was not demonstrated, and the cutoff was outcome-optimized, this exploratory analysis supports a clinically relevant LYVE1-associated stromal/myeloid niche rather than a validated macrophage-specific prognostic biomarker. To nominate regulatory features of the LYVE1-associated branch, transcription-factor activity analysis identified Yin Yang 1 (YY1) among the regulators enriched in this state (Fig. 6A). Consistent with reported links between YY1 and M2-like macrophage programs (33), ectopic YY1 expression in RAW264.7 macrophages increased Lyve1 and Mrc1 expression (Fig. 6B and C; Supplementary Fig. S6C). These data nominate YY1 as a candidate regulator and show that YY1 overexpression is sufficient to increase these markers in vitro; they do not establish YY1 necessity in vivo or indicate that DAC directly induces YY1 or LYVE1.

**Fig. 6.**
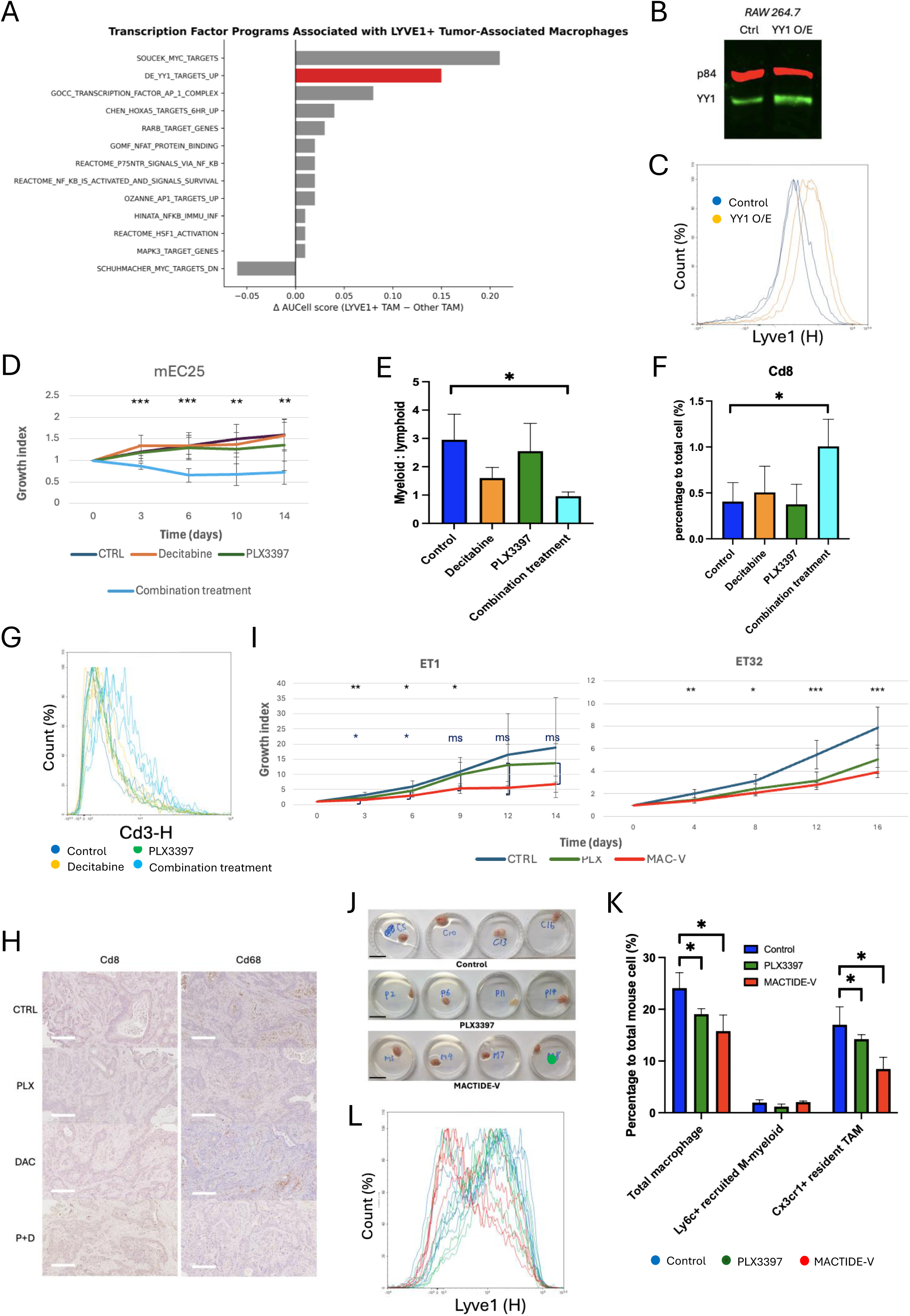
Immunocompetent efficacy and orthogonal targeting of Lyve1⁺ resident macrophages extend the therapeutic principle. (**A**) Transcription factor analysis of Lyve1⁺ resident tumor-associated macrophages (TAMs), showing enrichment of M2-associated transcriptional programs, including Yy1. (**B**) Immunoblot confirming YY1 overexpression in macrophages, with p84 as a loading control. (**C**) Relative Lyve1 expression in control and YY1-overexpressing (O/E) macrophages, demonstrating induction of Lyve1 following YY1 overexpression. (**D**) Tumor growth curves of mEC25 syngeneic tumors in C57BL/6 mice treated with vehicle, PLX3397, DAC, or the combination, showing superior antitumor efficacy and tumor regression with combination treatment in an immunocompetent setting. (**E**) Quantification (right) of the myeloid-to-lymphoid ratio following the indicated treatments, demonstrating a marked reduction after combination therapy. Gating: Macrophages as H-2Kd⁺Cd45⁺F4/80⁺. PMN cells as H-2Kd⁺Cd45⁺Ly6g⁺. Lymphoid cells were defined as H-2Kd⁺Cd45⁺Ly6c⁻Ly6g⁻. (**F**) Flow cytometric quantification of tumor-infiltrating CD8⁺ T cells, normalized to total murine cells, showing increased CD8⁺ T-cell infiltration following combination treatment. CD8⁺ T cells were gated as H-2Kd⁺Cd45⁺Ly6c⁻Ly6g⁻Cd3^+^Cd8^+^. (**G**) Relative Il2ra (Cd25) expression in tumor-infiltrating T cells across treatment groups, demonstrating enhanced T-cell activation following combination therapy. (**H**) Representative Cd8 immunohistochemistry and quantification confirming increased CD8⁺ T-cell infiltration in combination-treated tumors. Scale bar, 200 μm. (**I**) Tumor growth curves of ET1 (PLX3397-refractory) and ET32 (PLX3397-responsive) ESCC PDOX models treated with vehicle, PLX3397, or MACTIDE-V, demonstrating antitumor activity of MACTIDE-V in both models. (**J**) Representative images of ET1 PDOX tumors following the indicated treatments. (**K**) Flow cytometric analysis of resident-like and recruited-like TAM populations following vehicle, PLX3397, or MACTIDE-V treatment, demonstrating preferential depletion of resident-like TAMs by MACTIDE-V. (**L**) Flow cytometric quantification of Lyve1 expression in tumor TAMs, showing reduced Lyve1⁺ macrophages following MACTIDE-V treatment. *, P < 0.05; **, P < 0.01; ***, P < 0.001; ms, marginal significance (P < 0.1).

**Table 1.**
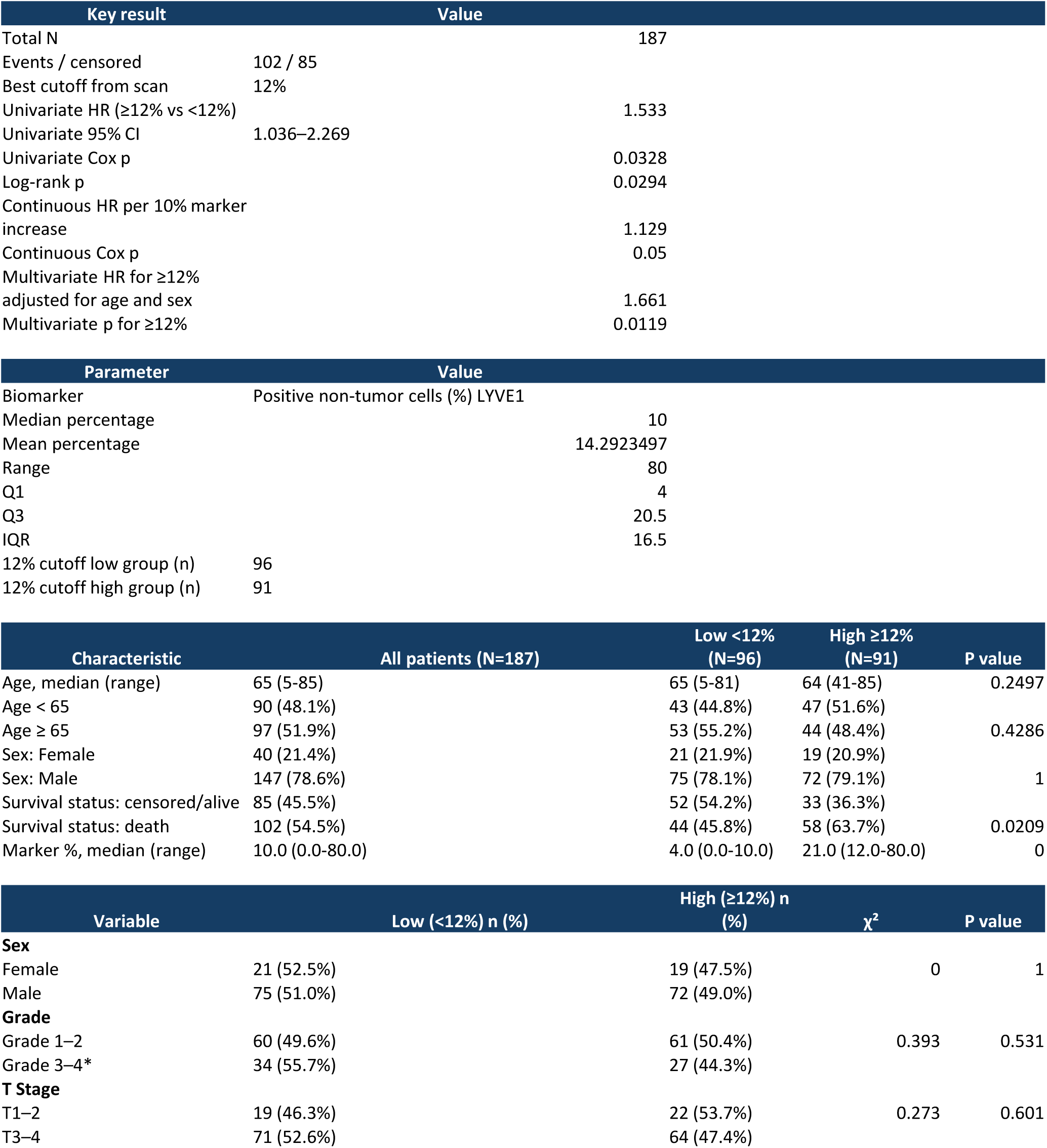
Summary of survival analyses for LYVE1 expression in an ESCC tissue microarray cohort.

### Immunocompetent and orthogonal perturbations extend the therapeutic principle

To determine whether dual-compartment control remained active in an intact immune system, we evaluated combined PLX3397 and DAC in the immunocompetent mEC25 ESCC model in C57BL/6 mice. The combination induced marked regression, whereas either monotherapy had limited activity; several combination-treated mice achieved near-complete regression (Fig. 6D; Supplementary Fig. S7A). Combination-treated tumors also showed a lower myeloid-to-lymphoid ratio, increased CD8-positive T-cell infiltration, and higher Cd25/Il2ra expression (Fig. 6E-H). In human ESCC single-cell data, macrophage LYVE1 expression inversely correlated with T-cell IL2RA expression (Supplementary Fig. S7B and S7C). These findings associate dual-compartment control with T-cell infiltration and activation markers, but do not establish T-cell-mediated regression.

As an orthogonal test of the tractability of the resident-like branch, we evaluated MACTIDE-V, a CD206-targeting peptide-drug conjugate developed to modulate CD206-expressing macrophages (34). MACTIDE-V suppressed tumor growth in two ESCC PDOX models, including ET1, which showed limited responsiveness to PLX3397 monotherapy (Fig. 6I and J). Treatment was not associated with significant changes in body weight or forelimb grip strength (Supplementary Fig. S8A and S8B). Flow cytometry showed a greater reduction in the resident-like than recruited-like compartment, accompanied by lower tumor Lyve1 expression (Fig. 6K and L). This independent perturbation provides preliminary proof of concept that the LYVE1/CD206-enriched branch is therapeutically tractable, although the cellular specificity and mechanism of MACTIDE-V in ESCC require further investigation.

## Discussion

Macrophage-directed therapy is commonly framed as a depletion problem. Our data identify a coverage problem: ESCC can maintain myeloid support through complementary treatment-defined persistence and replenishment vulnerabilities. CSF1R blockade reduced established TAMs but was followed by accumulation of Ly6C/CCR2-positive monocytic and Ly6G-positive granulocytic populations, whereas low-dose DAC preferentially restricted the recruited arm while leaving a LYVE1-associated macrophage state comparatively preserved. Combined treatment suppressed both blind spots and produced greater and more sustained control than either agent alone across PDOX, CDX, orthotopic, and immunocompetent models. The principal advance is therefore not simply the description of TAM heterogeneity or another combination regimen, but the resolution and joint disruption of complementary myeloid states that explain monotherapy escape.

Compensatory myeloid responses after CSF1R blockade have been described previously, including fibroblast-driven PMN-MDSC recruitment and recurrence supported by rebound macrophages (22, 24). Here, the compensatory response encompassed both monocytic and granulocytic compartments and was interpreted alongside reciprocal sparing of a LYVE1-associated state under DAC. The anti-Ly6G experiment separated early response from sustained control: Ly6G-positive cell depletion reproduced the initial regression achieved with PLX3397 plus DAC, but tumors subsequently regrew. Granulocytes therefore contribute to tumor maintenance after TAM depletion, yet neutrophil-directed depletion alone does not account for the sustained control achieved by broader restriction of the recruited arm. The actionable unit may therefore be the recruited escape arm rather than any single recruited cell type.

The reciprocal monotherapy responses define a pharmacologic division of vulnerability, not absolute cellular selectivity. PLX3397 reduced several CSF1R-dependent macrophage populations but did not prevent accumulation of Ccr2/Ly6c2/S100a9-expressing monocytic cells or Ly6G-positive granulocytes. Under the tested low-dose schedule, DAC strongly reduced these recruited populations while comparatively sparing a Lyve1/Mrc1-positive macrophage state. Decitabine can also alter tumor-cell epigenetics, antigen presentation, hematopoietic progenitors, stromal programs, and other immune lineages (25–29, 35). The absence of combination-specific organoid cytotoxicity supports a substantial in vivo microenvironmental contribution but does not assign efficacy exclusively to myeloid cells. Accordingly, the combination demonstrates complementary pharmacologic coverage without establishing formal synergy or a mechanism restricted to two cell types.

Single-cell profiling mapped this pharmacologic division onto treatment-resolved myeloid states. TAM-1, TAM-2, and TAM-3 formed a C1qa-positive continuum with progressive attenuation of inflammatory, interferon, antigen-presentation, and phagocytosis-associated programs. A C1qa-negative, Ccr2/Ly6c2-high inflammatory monocytic-like population remained distinct and was preferentially reduced by DAC, whereas the LYVE1-associated state persisted. Treatment response therefore provided an orthogonal criterion that added functional context to existing ESCC myeloid taxonomies: the contribution is not another static atlas, but a map of recognizable states onto complementary therapeutic vulnerabilities (32, 36). Flow cytometry and orthogonal perturbations further supported this organization, rather than clustering alone. Nevertheless, recruited-like and resident-like remain operational designations: marker expression, pseudotime position, and differential drug sensitivity do not establish developmental origin, and the C1qa-negative compartment may include inflammatory monocytes, immature macrophages, and M-MDSC-like cells (10–13, 16, 37–39).

Broad suppression of the treatment-resolved states coincided with changes beyond measured myeloid abundance. Combination-treated tumors contained fewer CAFs and endothelial cells, reduced CD34-positive vasculature, and increased antigen-processing, antigen-presentation, and lymphoid-associated programs. This pattern is consistent with disruption of a myeloid-supported stromal and vascular niche, particularly because LYVE1-positive macrophages have been linked to perivascular support, matrix organization, angiogenesis, and tumor growth in other tissues (14–18). In the mEC25 model, dual-compartment control also coincided with a lower myeloid-to-lymphoid ratio, increased CD8-positive infiltration, and higher Il2ra expression. However, these associations do not establish that loss of fibroblasts, vasculature, LYVE1-associated macrophages, or any lymphoid effector directly caused regression; tumor shrinkage, viable-tissue sampling, dissociation, and differential recovery may also influence apparent abundance. CD8- and NK-cell depletion studies will be required to define immune-effector dependence (40–43).

Human analyses supported disease relevance while preserving important interpretive boundaries. Reanalysis of a human single-cell dataset identified antigen-presenting, differentiated TAM, and LYVE1-associated macrophage programs corresponding to those observed in murine tumors, extending prior studies of ESCC myeloid and stromal heterogeneity (32, 36, 44). Treatment-induced IL34-associated CD163-positive TAM accumulation has also been reported in ESCC, reinforcing the view that myeloid composition is a dynamic determinant of therapeutic response rather than a static disease correlate (45). At the tissue level, high LYVE1 staining was associated with shorter overall survival. The convergence of controlled perturbation, cross-species transcriptional correspondence, and tissue-level outcome association supports disease relevance of the LYVE1-associated program, while not validating a macrophage-specific prognostic biomarker.

The translational implication is state-aware rather than bulk macrophage-directed therapy. Clinical studies of CSF1R-directed agents have demonstrated target engagement but limited activity in unselected solid tumors, consistent with the possibility that depletion of one macrophage compartment is insufficient when alternative states persist or emerge (43). Pharmacodynamic assessment could therefore incorporate multiparameter signatures of the C1qa-positive TAM continuum, CCR2/Ly6C-positive recruited cells, and the LYVE1/MRC1-associated tissue-supportive state rather than relying on total CD68- or F4/80-positive abundance alone. Pexidartinib and decitabine provide proof-of-principle perturbations, not a proposed clinical regimen; their pleiotropic effects, scheduling, hematologic consequences, and tolerability require careful evaluation. The anti-Ly6G comparison cautions that narrow neutrophil targeting may not reproduce broader recruited-arm control, whereas MACTIDE-V provides orthogonal, preliminary evidence that a CD206-enriched resident-like compartment can also be perturbed (34). Together, these findings motivate biomarker-guided, multi-axis myeloid strategies—potentially combined with checkpoint blockade—rather than empirical bulk TAM depletion. Whether this persistence-replenishment framework extends to other macrophage-rich squamous tumors requires direct testing.

Several limitations define the scope of this framework. Fate mapping, parabiosis, bone marrow chimerism, or lineage barcoding will be required to distinguish resident maintenance from monocyte-derived replacement and to resolve interconversion among treatment-defined states. The mechanism by which low-dose DAC reduces recruited myeloid populations remains unresolved across marrow, blood, and tumor compartments, including possible effects on progenitor production, survival, trafficking, and differentiation. Lineage-specific perturbations are needed to define the contributions of Ccr2-positive monocytic cells, Ly6G-positive granulocytes, and the LYVE1-associated macrophage state, while CD8- and NK-cell depletion will be required to identify immune effectors of regression. Spatial profiling and multiplex colocalization are needed to establish the localization and cellular specificity of the LYVE1-associated niche in human ESCC. Biological-replicate-level differential-abundance testing, complete trajectory and cross-species mapping methods, and clinically relevant exposure, scheduling, toxicity, and reversibility of broad myeloid suppression will also be essential for translation.

Together, these data shift the therapeutic target from bulk macrophage depletion to state-aware coverage of complementary myeloid vulnerabilities. In ESCC, a C1qa- positive TAM continuum, a recruited Ccr2/Ly6c2-high inflammatory monocytic-like state, Ly6G-positive granulocytic populations, and a LYVE1-associated tissue-supportive state showed non-equivalent treatment sensitivities that explained monotherapy escape and combination response. By coupling defined states to cross-model phenotyping, human correspondence, and orthogonal perturbation, this persistence-replenishment framework provides a testable basis for biomarker-guided myeloid therapy.

## Acknowledgments

This work was supported by the Health and Medical Research Fund of the Health Bureau, Hong Kong SAR Government (06171566 and 11221936 to V.Z.Y.); the Research Grants Council (Theme-based Research Scheme T12-701/17-R to M.L.L. and S.L.); and the Spanish State Research Agency (Agencia Estatal de Investigación, AEI; PDC2025-165219-I00 to P.S.). We thank the Leibniz Institute DSMZ–German Collection of Microorganisms and Cell Cultures GmbH for providing the KYSE cell lines. We thank the Centre for Comparative Medicine Research, Li Ka Shing Faculty of Medicine, The University of Hong Kong, for animal facilities, and the Centre for PanorOmic Sciences for sequencing and imaging support.

## Methods

### Chemical reagents

All small-molecule compounds were purchased from MedChemExpress (Monmouth Junction, NJ), unless otherwise specified. MACTIDE-V was synthesized by the Proteomics Service of CNB-CSIC (Madrid, Spain).

### Animals

Female BALB/cAnN-nu or C57BL/6 mice older than 7 weeks were used. All procedures were approved by the Committee on the Use of Live Animals in Teaching and Research and conducted in the AAALAC International-accredited Centre for Comparative Medicine Research, Li Ka Shing Faculty of Medicine, The University of Hong Kong, under licenses issued by the Hong Kong SAR Government Department of Health.

### Tumor models and treatment

PDOX and CDX models of ESCC were generated subcutaneously or orthotopically as described previously (46–48). Immunocompetent experiments used the mEC25 model in C57BL/6 mice (49). When tumors reached 150 to 200 mm3, mice were randomized to treatment groups. PLX3397 was administered by oral gavage at 40 mg/kg daily, and DAC was administered intraperitoneally at 1 mg/kg every 3 days. Combination treatment used concurrent administration; control mice received the corresponding vehicles. Anti-Ly6G antibody was administered intraperitoneally according to the schedule shown in Fig. 2H. Group sizes ranged from 6 to 9 animals, with most experiments using 8 or 9 mice per group. Tumors were measured two to three times weekly, and mice were monitored to humane endpoints. Tumor growth was expressed as a growth index calculated by dividing tumor volume on each measurement day by tumor volume at treatment initiation.

### In vivo imaging

Orthotopic CDX models generated with luciferase-labeled ESCC cells were monitored by bioluminescence imaging using the IVIS Lumina X5 system (PerkinElmer) as described previously (47). D-luciferin was administered intraperitoneally at 150 mg/kg 7 minutes before imaging. Three-dimensional tumor signals were captured using the Xenogen IVIS Spectrum. Treatment began when the signal exceeded 1 × 10^6 photons per second. Tumor growth was expressed as a growth index calculated by dividing the bioluminescence signal within the tumor region of interest on each imaging day by the corresponding signal at treatment initiation.

### Blood and tissue collection

Mice were euthanized by ketamine-xylazine overdose at the indicated endpoints. Blood was centrifuged at 2,000 × g for 10 minutes to obtain plasma. Tumors were fixed in 10% neutral-buffered formalin, enzymatically dissociated into single-cell suspensions, or preserved in RNAlater for molecular analyses.

### Histology and immunohistochemistry

Formalin-fixed, paraffin-embedded tissues were analyzed by immunohistochemistry using standard protocols. Primary antibodies are listed in Supplementary Table S2. Signals were visualized with horseradish peroxidase-conjugated secondary antibodies and 3,3′-diaminobenzidine, followed by hematoxylin counterstaining. Quantification used the numbers of tumors and fields specified in the figure legends.

### Flow cytometry

Single-cell suspensions were generated from tumors by enzymatic dissociation as described previously (46). Antibody panels are listed in Supplementary Table S2. Analyses included total leukocytes, F4/80-positive macrophages, Ly6C/CX3CR1-defined macrophage-enriched compartments, Ly6G-positive cells, NK cells, and lymphoid populations. Unless otherwise stated, immune populations were normalized to total murine cells to facilitate comparison across models and implantation sites.

### RNA extraction and quantitative PCR

RNA was isolated from RNAlater-preserved tissues using TRIzol (Thermo Fisher Scientific). Complementary DNA was synthesized from 0.2 μg RNA using the QuantiNova Reverse Transcription Kit (Qiagen). Quantitative reverse-transcription PCR was performed on a Roche LightCycler 480 II system. Primer sequences are provided in Supplementary Table S3.

### Bulk RNA sequencing and analysis

Poly(A)-enriched bulk RNA sequencing was performed on PDOX tumors treated with PLX3397 or vehicle at the HKUMed Centre for PanorOmic Sciences Genomics and Bioinformatics Cores as described previously (46). Reads were aligned separately to the human hg38 and mouse mm10 transcriptomes. Differential-expression analysis and downstream visualization were performed in Partek Flow Genomic Analysis Software (Partek Inc., St. Louis, MO).

### Single-cell RNA sequencing and analysis

PDOX tumors were enzymatically dissociated, and viable single-cell suspensions were processed using the Chromium Single Cell 5′ platform (10x Genomics, Pleasanton, CA). Libraries were sequenced at the HKUMed Centre for PanorOmic Sciences Genomics and Bioinformatics Cores. Cell Ranger version 6.0.0 was used with a combined hg38-mm10 reference genome. Quality control, normalization, principal-component analysis, Louvain clustering, and marker identification were performed in Partek Flow. Cell numbers and cluster-defining markers are listed in Supplementary Table S4.

Gene-set enrichment analysis was performed in R version 4.3.3 using clusterProfiler with Hallmark and Gene Ontology Biological Process gene sets. Gene sets with P < 0.05 in the source analysis were considered significant. AUCell was used to calculate gene-set activity scores.

### Public human ESCC single-cell analysis

A publicly available ESCC single-cell RNA-sequencing dataset was reanalyzed in R using a workflow aligned with the murine analysis (32). Monocytes and dendritic cells were excluded before macrophage reclustering, and cross-species comparisons were used to relate human and murine TAM states.

### Human ESCC tissue microarrays and survival analysis

LYVE1 immunohistochemical staining was evaluated in two ESCC tissue microarrays (HEso-Squ172Sur-01 and HEso-Squ180Sur-03) comprising 187 patients. Survival analyses used the combined cohort without stratification by array set. Overall survival was defined with death as the event and survival at last follow-up as censoring. Patients were dichotomized as LYVE1-high (≥12% positive staining) or LYVE1-low (<12%) using an outcome-optimized cutoff. Survival was evaluated by Kaplan-Meier analysis and log-rank testing. Hazard ratios were estimated using univariable and multivariable Cox proportional-hazards models; the multivariable model adjusted for age and sex. Additional clinicopathologic and statistical details are provided in Supplementary Table S1.

### MACTIDE-V synthesis and treatment

MACTIDE-V was provided by Pablo Scodeller and synthesized by the Proteomics Service of CNB-CSIC (Madrid, Spain). The conjugate was Vert-Ahx-G(CTKSIPPIC)SPGAK-OH, where Vert denotes verteporfin, Ahx denotes aminohexanoic acid, the cysteines form a disulfide bond, and the peptide has a free C-terminus. Detailed synthesis and characterization are described in ref. 34. MACTIDE-V was dissolved in PBS and administered intraperitoneally at 3.4 mg/kg every other day.

### Organoid viability and YY1 overexpression

ESCC organoids were established in-house as described previously (46). Organoids were treated with PLX3397, DAC, or the combination to evaluate direct treatment-associated cytotoxicity. For YY1 studies, murine *Yy1* cDNA was ectopically expressed in RAW264.7 macrophages using a lentiviral system as described previously (50). The *Yy1* coding sequence was cloned into a lentiviral expression vector, followed by lentiviral production and transduction. *Lyve1* and *Mrc1* expression were assessed after YY1 overexpression.

### Statistical analysis

Quantitative data are presented as means with 95% confidence intervals unless otherwise stated. Tumor-growth curves were compared using two-way repeated-measures ANOVA. Two-group endpoint comparisons used unpaired, two-tailed Student t tests. Survival was analyzed as described above. All tests were two-sided, and P < 0.05 was considered statistically significant. Analyses were conducted using Excel, GraphPad Prism 9, and R.

---

**Supplementary Fig. S1.**
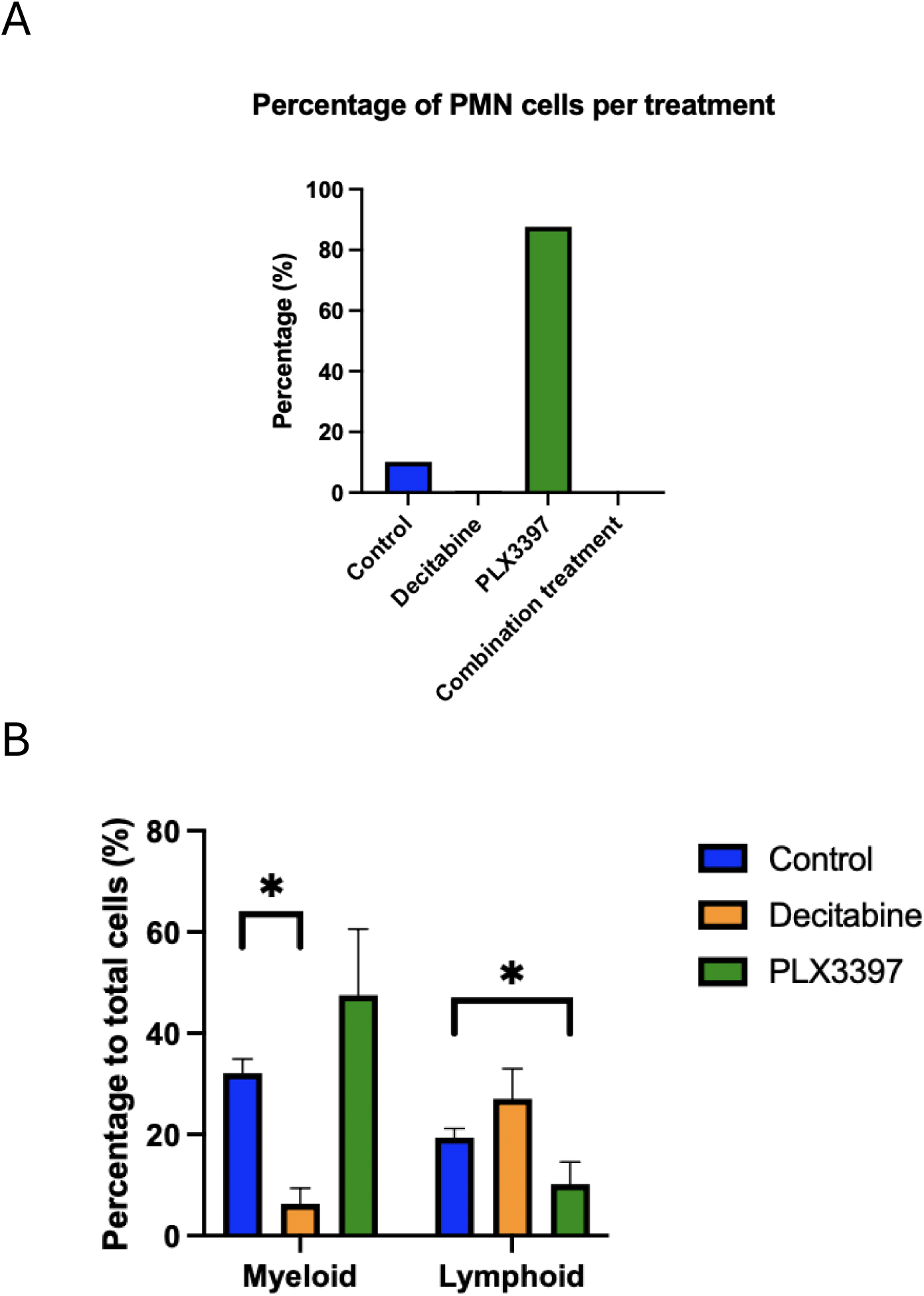
Decitabine preferentially depletes myeloid cells while sparing lymphoid populations. **(A)** Flow cytometric quantification of tumor-infiltrating polymorphonuclear (PMN) cells in PDOX tumors following the indicated treatments. **(B)** Summary of flow cytometric analysis showing that decitabine (DAC) selectively reduced myeloid cells while largely sparing lymphoid cells. Gating strategy: lymphoid cells, H-2Kᵈ⁺CD45⁺ cells with low FSC/SSC; myeloid cells, H-2Kᵈ⁺CD45⁺CD11b⁺; PMNs, H-2Kᵈ⁺CD45⁺Ly6G⁺. Data are presented as mean ± SEM. *P < 0.05.

**Supplementary Fig. S2.**
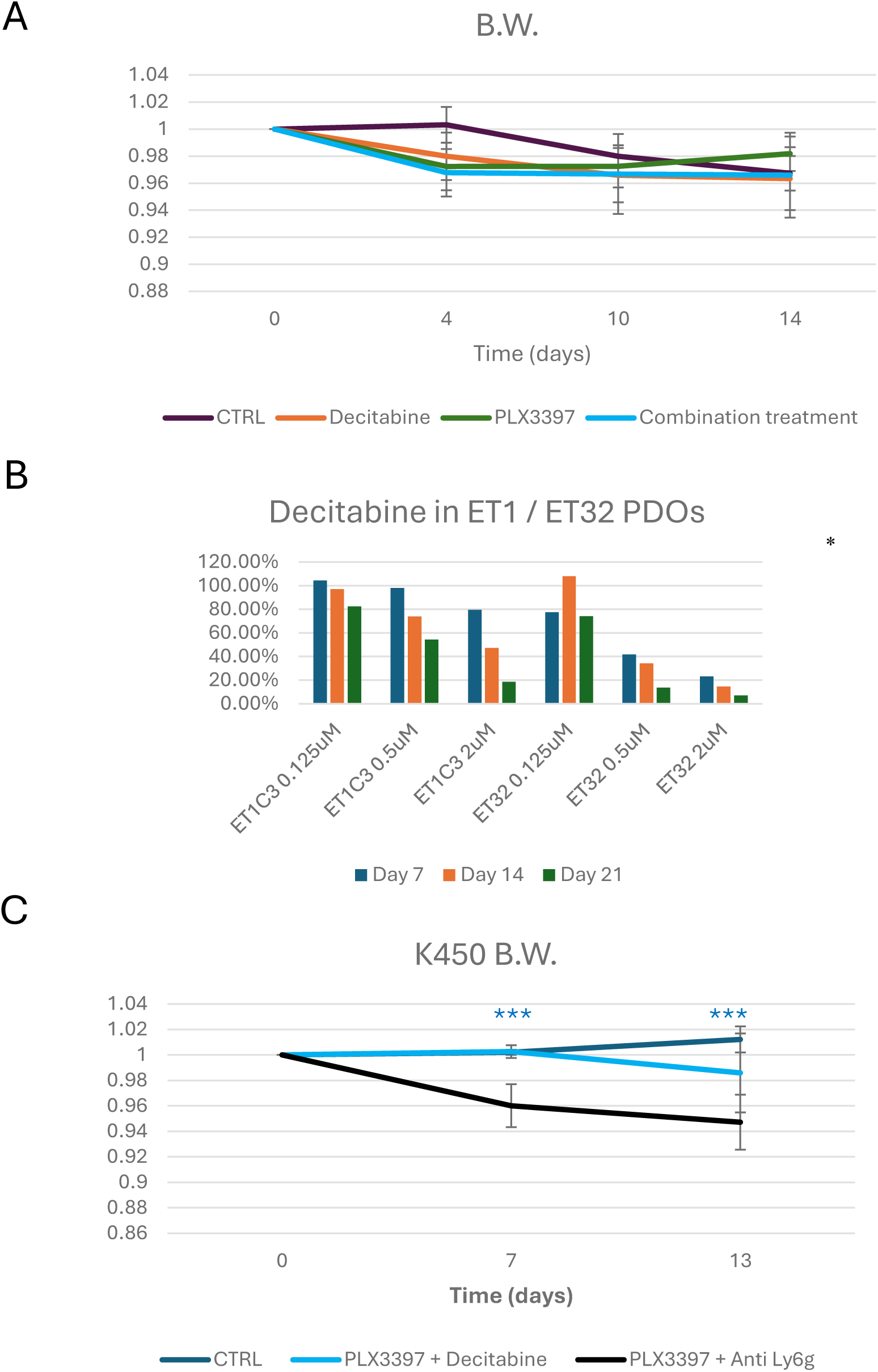
Combination therapy depletes CD206⁺ macrophages without increasing treatment-related toxicity. **(A)** Body weight changes of PDOX-bearing mice treated with the indicated monotherapies, showing no significant treatment-related weight loss. **(B)** Cell viability of ET1 and ET32 patient-derived organoids (PDOs) following treatment with increasing concentrations of DAC, measured by luminescence-based cell viability assay. (C) Body weight changes of CDX-bearing mice treated with the indicated combination therapies. Showing the PLX3397 + Anti-Ly6g combination treatment resulted in lowered body weight.

**Supplementary Fig. S3.**
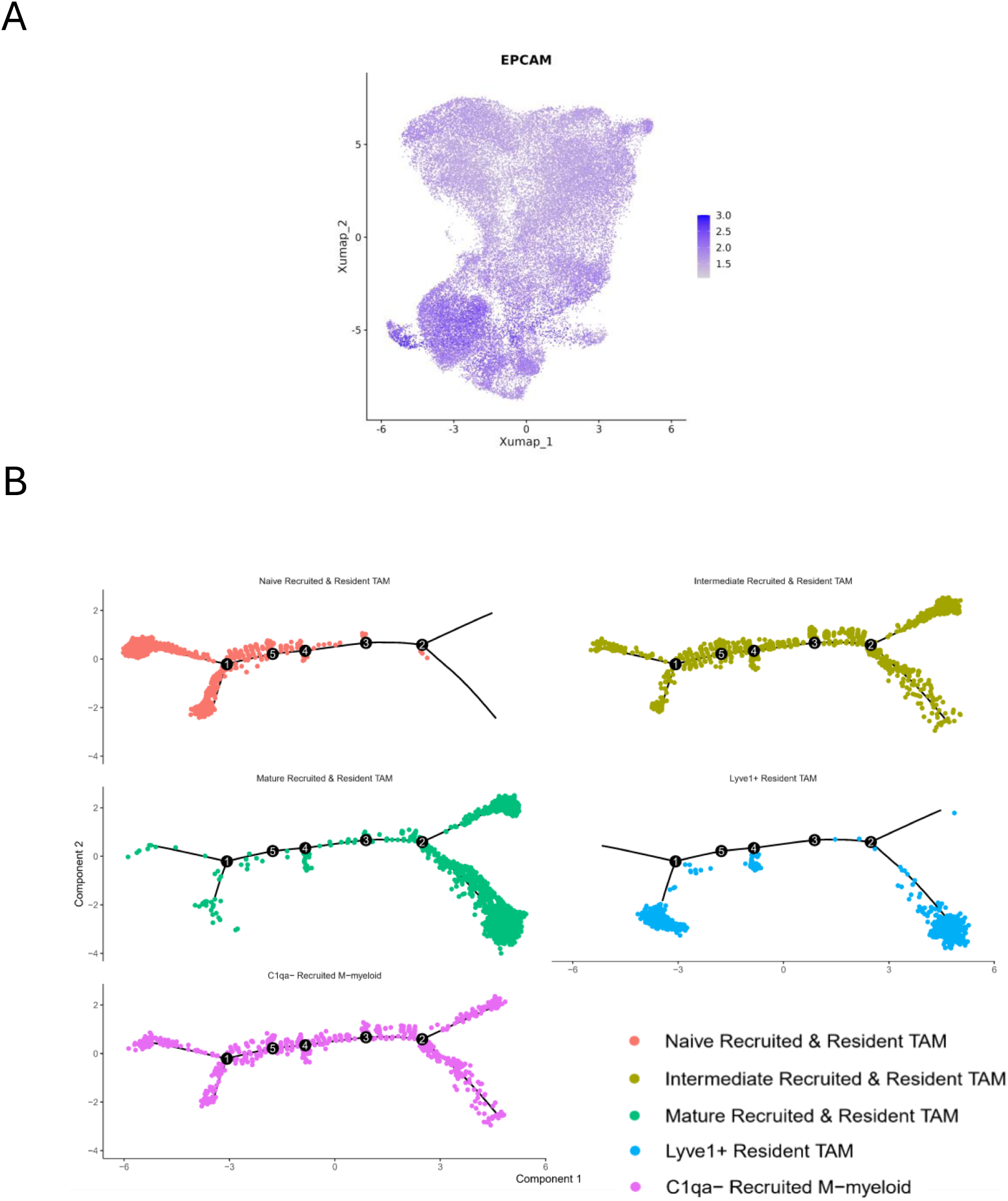
Single-cell characterization of human ESCC cells and macrophage differentiation trajectory. **(A)** UMAP of human ESCC tumor cells (EPCAM⁺) identified from the integrated single-cell RNA-sequencing dataset. **(B)** Pseudotime trajectory analysis of the five M-myeloid cell clusters, illustrating the inferred differentiation continuum from naïve to mature tumor-associated macrophage (TAM) states, including the C1qa^-^ recruited M-myeloid and LYVE1⁺ resident TAM population.

**Supplementary Fig. S4.**
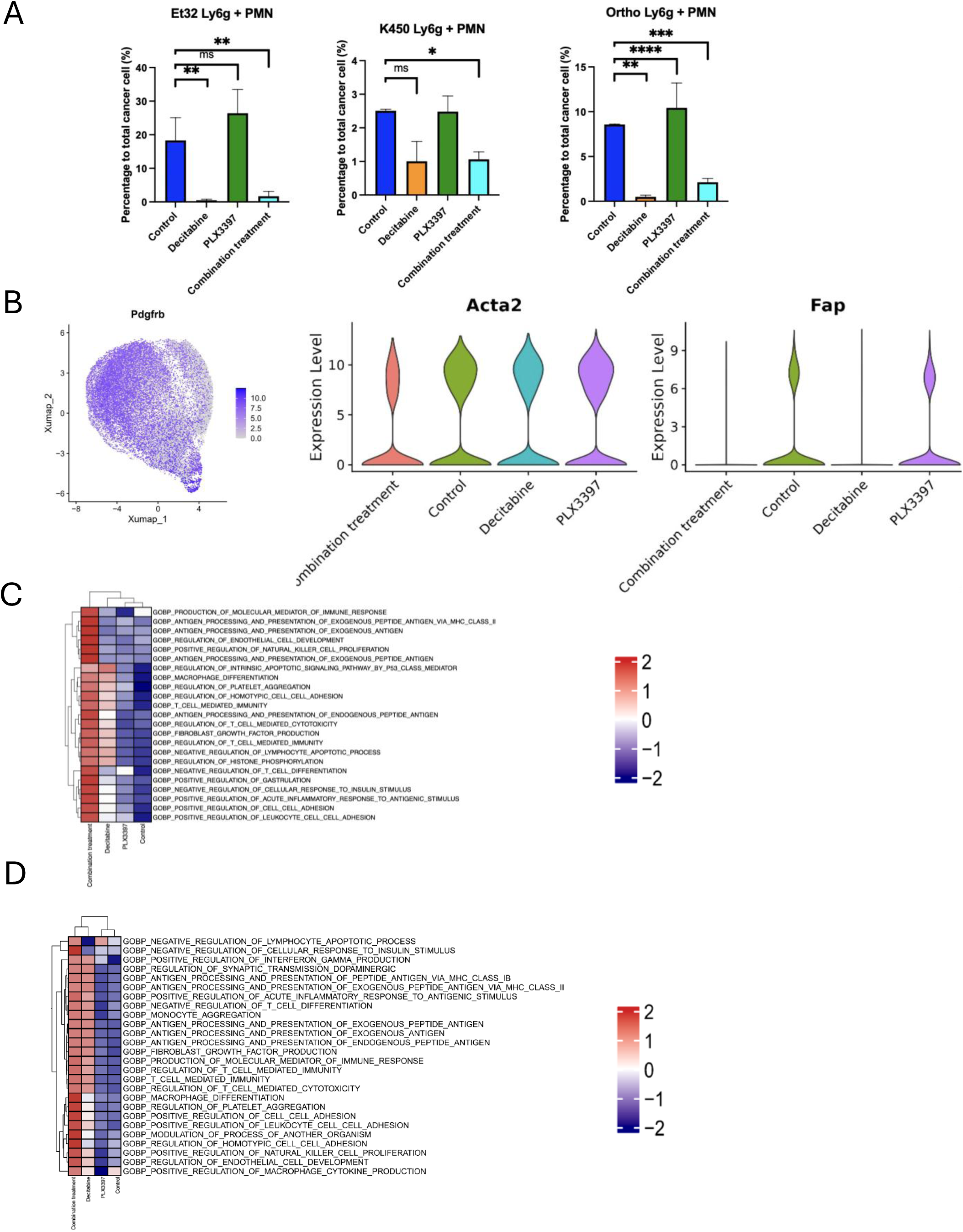
Additional analyses of stromal remodeling following dual myeloid targeting. **(A)** Flow cytometric quantification of tumor-infiltrating polymorphonuclear (PMN) cells across treatment groups, normalized to total murine cells. PMNs were gated as H-2kd⁺Cd45⁺Ly6g⁺ cells. **(B)** UMAP of cancer-associated fibroblasts (CAFs) with *Pdgfb* expression highlighted (left) and expression levels of *Acta2* and *Fap* across treatment groups (right). **(C)** Gene Ontology Biological Process (GOBP) enrichment analysis of CAFs following the indicated treatments. **(D)** GOBP enrichment analysis of ESCC cells following the indicated treatments.

**Supplementary Fig. S5.**
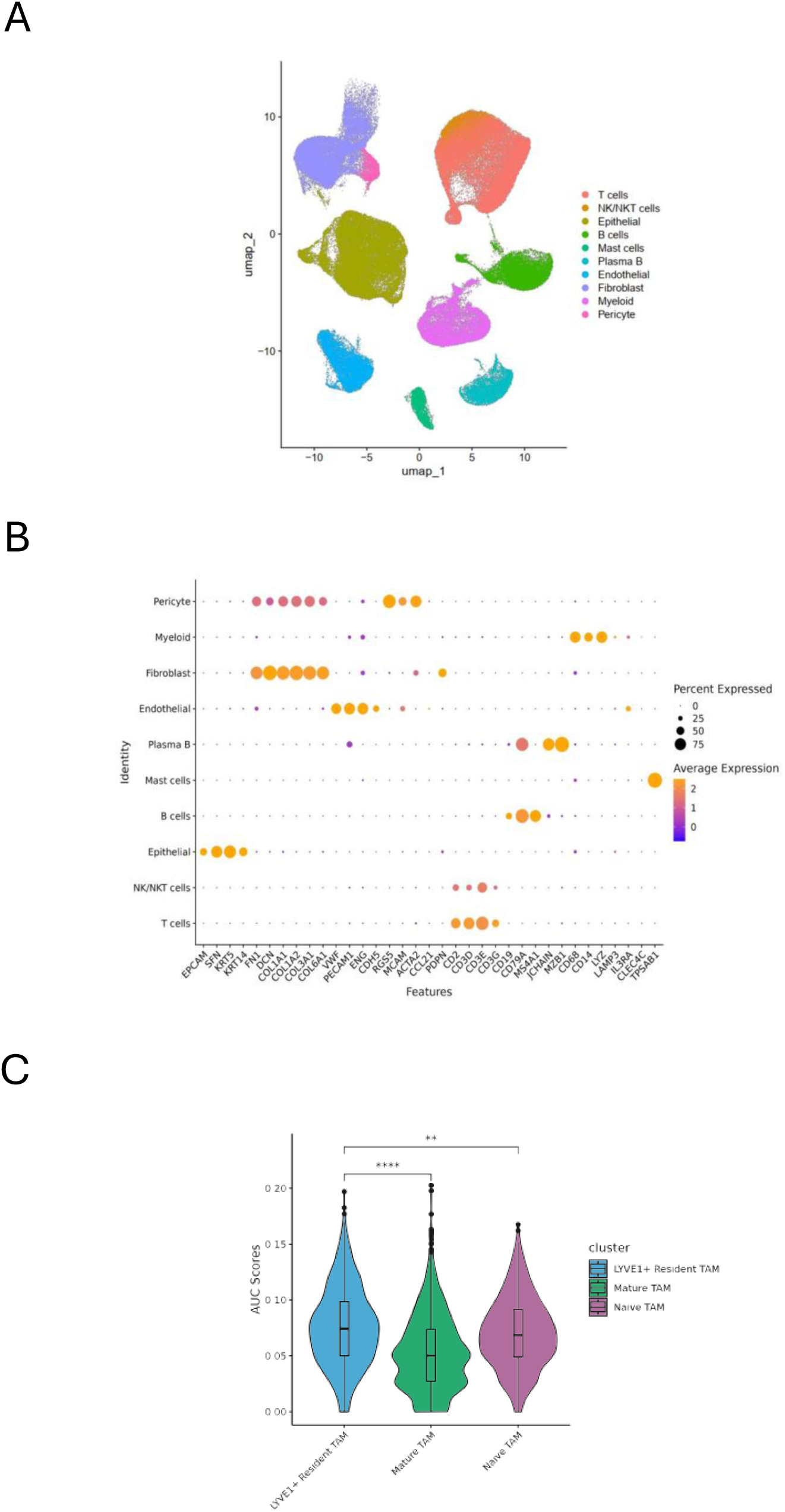
Single-cell transcriptomic annotation of human ESCC. **(A)** UMAP of all cells from the human ESCC single-cell RNA-sequencing dataset, showing unsupervised clustering of the major cellular populations. (**B)** Dot plot of canonical marker genes used to define each annotated cell type.**(C)** Hallmark pathway enrichment analysis demonstrating preferential activation of angiogenesis in the murine Lyve1⁺ resident TAM

**Supplementary Fig. S6.**
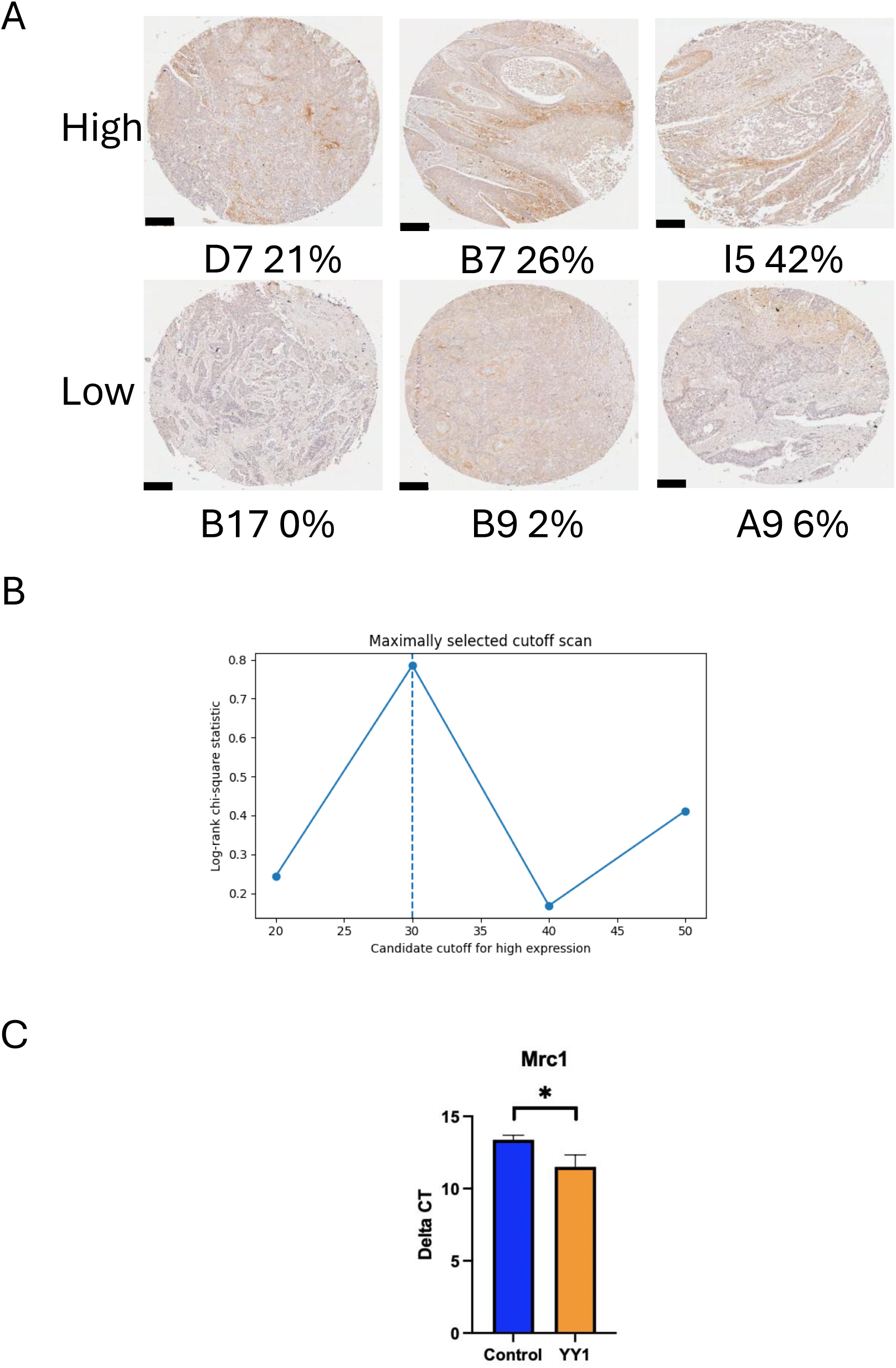
Validation of LYVE1 clinical scoring and YY1-mediated macrophage polarization. **(A)** Representative images illustrating LYVE1 immunohistochemical scoring in the ESCC tissue microarray and derivation of the outcome-optimized 12% cutoff for classification into LYVE1-low and LYVE1-high tumors. Scale bar, 200 µm. **(B)** Maximally selected rank statistic identifying the optimal LYVE1-positive cell cutoff (12%) for overall survival analysis. **(C)** Relative *Mrc1* expression in control and YY1-overexpressing macrophages measured by quantitative PCR, demonstrating increased *Mrc1* expression following YY1 overexpression.

**Supplementary Fig. S7.**
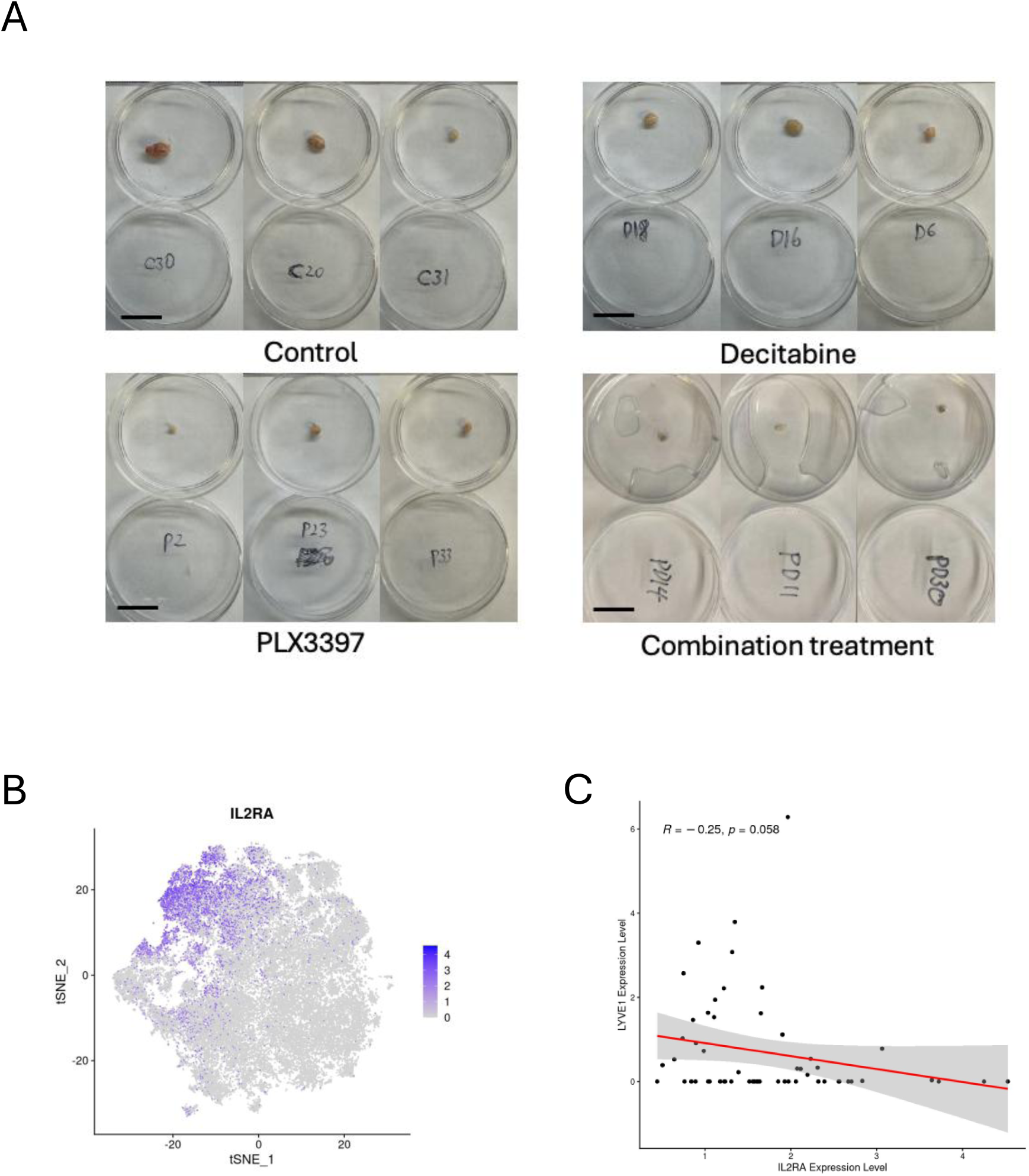
Additional analyses of combination therapy and human T-cell activation. **(A)** Representative images of mEC25 syngeneic tumors following vehicle, PLX3397, decitabine (DAC), or combination treatment, demonstrating marked tumor regression with combination therapy. **(B)** UMAP of tumor-infiltrating T cells from the human ESCC single-cell RNA-sequencing dataset (n = 60 tumors), with *IL2RA* expression highlighted. **(C)** Correlation between macrophage *LYVE1* and T-cell *IL2RA* expression in human ESCC single-cell data, demonstrating an inverse association (Spearman correlation; *P* = 0.058).

**Supplementary Fig. S8.**
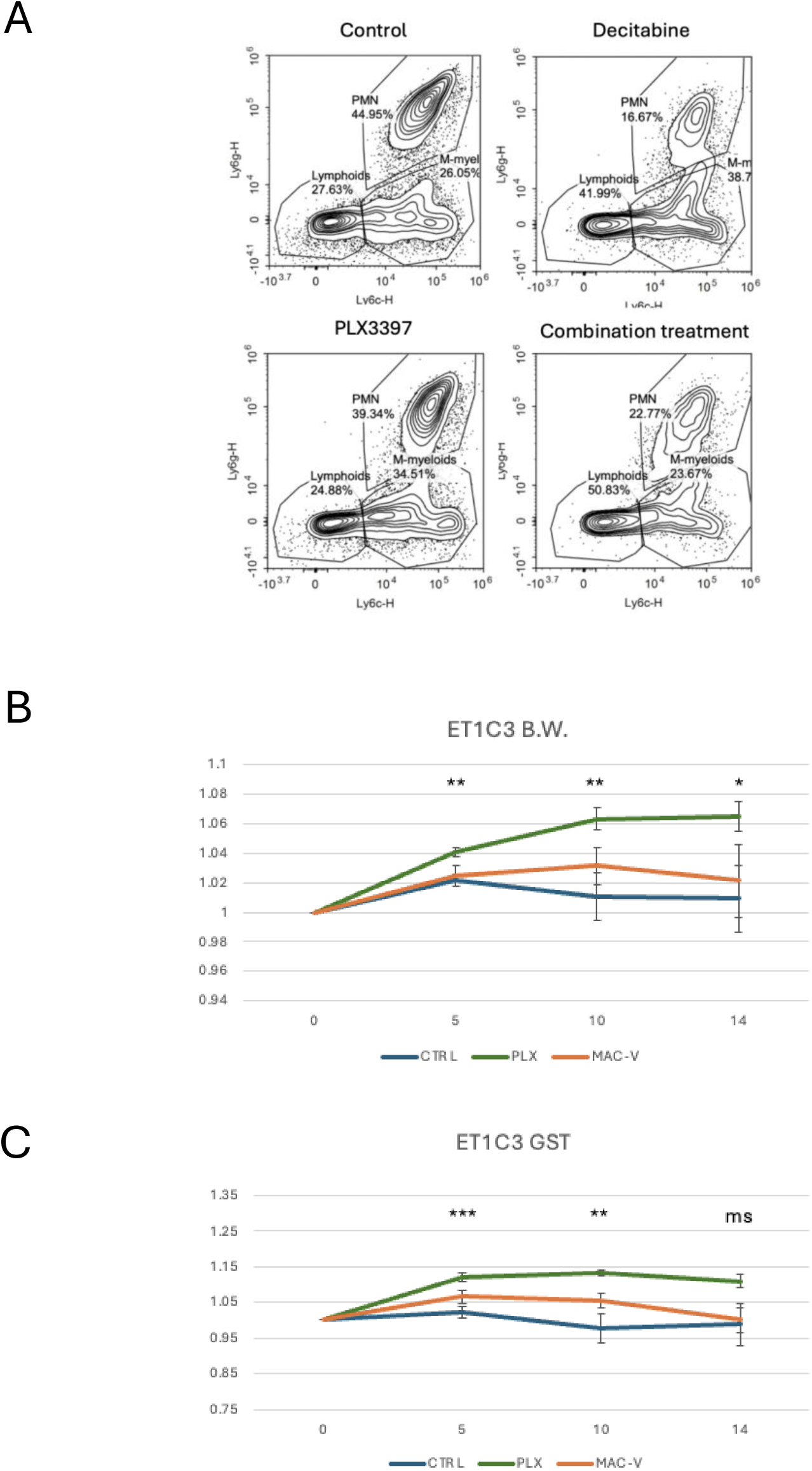
MACTIDE-V is well tolerated in vivo. **(A)** Body weight changes of mice treated with the indicated therapies. **(B)** Forelimb grip strength measurements following treatment, demonstrating preserved muscle function and tolerability of MACTIDE-V. **(C)** Representative flow cytometry plots showing M-myeloid, PMN, and lymphoid cell populations across treatment groups.

## Notes

### Competing Interest Statement

The authors have declared no competing interest.

